# Accumulation of PD-1+ TIGIT+ T cells in the liver after local antigen reactivity and during autoimmune hepatitis

**DOI:** 10.1101/2025.08.29.672936

**Authors:** Thomas Guinebretière, Anaïs Cardon, Jean-Paul Judor, Pierre-Jean Gavlovsky, Fanny Roux, Virginie Huchet, Caroline Chevalier, Marion Khaldi, Edouard Bardou-Jacquet, Laure Elkrief, Adrien Lannes, Christine Silvain, Matthieu Schnee, Florence Tanne, Sara Lemoinne, Eleonora De Martin, Fabienne Vavasseur, Arnaud Nicot, Sophie Brouard, Jean-François Mosnier, Jérôme Gournay, Sophie Conchon, Amédée Renand

## Abstract

In autoimmune hepatitis (AIH), hepatocellular damage is linked to an accumulation of autoreactive T cells in the liver of patients, but how these cells emerge in the tissue remains unclear. Here we used a mouse model based on recombination-dependent inducible expression of influenza A hemagglutinin (HA) by hepatocytes to investigate initiation of liver antigen-specific response. Our study revealed that peripheral immunization, unlike inflammatory triggers, is essential to initiate an immune response against a liver antigen. We showed that liver T cell reactivity after peripheral immunization is marked by PD-1 and TIGIT co-expression and that the frequency of PD-1^+^ TIGIT^+^ HLA-DR^+^ CD38^+^ CD8 T cells in the blood of AIH patients is associated with liver injury. Our findings suggest a potential influence of the peripheral immunization for the liver-antigen-specific responses during AIH. Liver tissue-activated T cells probably recirculate during active phase of the disease, unveiling potential immunomarkers to monitor disease activity.

**Highlights:**

- Peripheral immunization rather than local inflammation induces an immune response against a hepatic antigen
- PD-1 and TIGIT co-expression by T cells is found after tissue antigen-specific T cell reactivity
- Frequency of circulating PD-1^+^ TIGIT^+^ HLA-DR^+^ CD38^+^ CD8 T cells is associated with AIH disease activity

**In brief:** Guinebretière et al. demonstrate that after peripheral immunization, liver-antigen-specific T cell accumulation in the tissue is marked by local PD-1/TIGIT co-expression, a phenotype shared with liver and circulating T cell subsets enriched in active autoimmune hepatitis (AIH) patients. These findings suggest the influence of peripheral immunization on the initiation of AIH and provide potential immunomarker of AIH activity.

## Introduction

In healthy condition, antigen presentation is finely regulated in the liver to limit inflammation and preserve tissue functions. In autoimmune hepatitis (AIH), a rare chronic inflammatory disease of the liver, hepatic tolerance is chronically impaired, which leads to local immune cell infiltrates and progressive destruction of hepatocytes. Liver self-antigens such as SEPSECS, CYP2D6 and FTCD are targeted in the disease and specific autoantibodies and autoreactive CD4 T cells are detectable in the blood of AIH patients^1–4^. Both autoantibody production and autoreactive CD4 T cell detection in the blood highlight the complete adaptive immune response against the liver-self-antigens^2–4^. Recently, the transcriptomic signature of circulating autoreactive CD4 T cells from patients has been deeply characterized at the single-cell level^2,3^. This revealed a B-helper and immuno-exhausted signature with immune checkpoint upregulation, showing similarities with the profile of tissue autoreactive T cells in other autoimmune diseases and with chronically activated tissue-resident memory (Trm)-like cells during cancer and virus infection^2,3,5–14^. Furthermore, TCR clonotype analysis and comparison between the liver and the blood of patients suggest that circulating autoreactive CD4 T cells derived from the liver where they had acquired their transcriptomic profile by reacting in the tissue^3^. However, the origin of these autoreactive T cells is still unknown and the late diagnosis of AIH patients prevents the study of disease onset from human biological materials.

Mouse models have been designed to reproduce AIH development. Most of them focused on the induction of an immune response against AIH-targeted liver self-antigens to strengthen the relevance to human disease. Thereby, intravenous injection of a hepatotropic adenovirus that codes for human self-antigens CYP2D6 or FTCD in mice resulted in chronic liver inflammation, fibrosis and production of autoantibodies^15–18^. This highlights how ‘molecular mimicry’, the cross-reactivity of the immune system between self-antigens and non-self-antigens delivered by a pathogen, can be involved in autoimmune response initiation^19^. However, the study of the initiation and dynamics of the antigen-specific response is limited in this context. While the spleen seems dispensable to initiate this autoreactivity in this model^20^, the need of priming of antigen-specific cells occurring in the periphery cannot be ruled out. In this way, intramuscular immunization against human CYP2D6 and FTCD leads to marked portal infiltrates and specific autoantibody production, suggesting this time that peripheral priming is enough to induce liver autoreactivity^21^. Otherwise, further signals seem to be involved. On the one hand, genetic background and susceptibility to autoimmune disorders have a significant influence in the previous models^15–18^. On the other hand, the importance of liver inflammatory triggers and tissue injury is more elusive. While long term hepatic inflammation is observed following mere immunization against a liver antigen^21,22^, viral or chemical triggers are also reported as essential bystanders for chronic liver autoreactivity induction^16,23^.

These works open interesting insights into how chronic hepatic inflammation targeting liver self-antigens is initiated, but investigating the origins of autoreactive T cells is technically challenging. Thus, TCR transgenic strategy enables the tracking of T cells specific for antigens expressed in the liver. In this way, spontaneous and chronic autoreactive hepatitis is observed in TCR transgenic mice with lymphocytic choriomeningitis virus glycoprotein (GP)-specific CD4 T cells or influenza virus hemagglutinin (HA)-specific CD8 T cells, when GP or HA are constitutively expressed in the liver, respectively^24,25^. Conversely, induction of liver HA expression in inflammatory condition is not associated with long term hepatic inflammation in TCR transgenic mice with HA-specific T cells^26^. However, the early initiation steps, such as peripheral immunization or local inflammation, have not been characterized in depth in these models where all CD4 and/or CD8 T cells are potentially pathogenic, which makes their relevance questionable when compared to the very low frequency of autoreactive T cells measured in AIH patients^2–4^.

Our group has previously developed a non-TCR transgenic mouse model based on the inducible expression of the model antigen HA by hepatocytes to study the liver immuno-regulation of antigen-specific T cells^3^. As reported, tamoxifen-induced HA expression by hepatocytes in HA immunized mice leads to the accumulation of HA-specific CD4 T cells in the liver that share gene expression signature with autoreactive CD4 T cells found in the blood of AIH patients^3^. Here, we used this mouse model to decipher the events needed to initiate a hepatocyte-antigen-specific response. We observed that the liver tolerance of HA withstands local danger and inflammatory triggers. We showed that peripheral immunization is an essential step to induce hepatocyte-antigen-specific T cell response. We found that hepatocyte-antigen-specific T cells accumulate in the tissue after peripheral immunization and that liver T cell reactivity is marked with PD-1 and TIGIT co-expression in both the mouse model and one AIH patient, making them distinctive markers of local antigenic reactivity in the liver tissue. In the blood of AIH patients, the frequency of circulating PD-1^+^ TIGIT^+^ HLA-DR^+^ CD38^+^ CD8 T cell subset was high and was associated with liver injury. These findings revealed a key connection between the liver and the periphery in hepatocyte-antigen-specific responses, with the importance of peripheral immunization to modulate the accumulation of PD-1^+^ TIGIT^+^ T cells in the liver and the potential recirculation of tissue-activated autoreactive T cells during chronic phase of AIH. Thus, tracking PD-1^+^ TIGIT^+^ HLA-DR^+^ CD38^+^ T cells in the blood of patients may be a source for studying T cell dependent-tissue-damages in AIH.

## Results

### Local inflammatory triggers are not sufficient to initiate adaptive immune response against HA expressed by hepatocytes

We have used a mouse model allowing the inducible expression of the hemagglutinin (HA) model antigen by hepatocytes^3^. Briefly, HA gene expression at the Rosa26 locus is restricted by LoxP sequences in *Rosa26^tm(HA)1Libl^* (RosaHAfl) mice, and inducible Cre recombinase expression depends on the hepatocyte-specific transthyretin (TTR) promotor in *TTR-Cre-ind* mice. Cross-breeding these strains resulted in *Rosa26^tm(HA)1Libl^/TTR-Cre-ind* (HA/iCre) mice, in which hepatocyte HA expression is induced and tolerated after tamoxifen treatment^3,27,28^ (Figure 1A). We asked whether adding local inflammatory triggers could break the hepatic tolerance to HA. The liver inflammation and anti-HA antibody production were compared in HA/iCre mice treated with tamoxifen alone or combined with two distinct inflammatory signals: an intravenous injection (i.v.) of a hepatotropic adenovirus vector (Ad Ct i.v.) or intraperitoneal injections of D-galactosamine hydrochloride (D-GalN) and lipopolysaccharide (LPS) (Figure 1B). Here, we compare liver inflammation induced by a virus with a tropism for hepatocytes (Ad Ct)^29^, and anti-bacterial-associated inflammation combined with hepatotoxicity (D-GalN/LPS)^30^. Histological analysis showed that tamoxifen-treated mice had normal liver parenchyma without inflammatory events (Figure 1C). After Ad Ct i.v. injection, tamoxifen-treated HA/iCre mice displayed marked infiltration of portal tracts by immune cells and high levels of transaminase in the serum, reflecting local adenoviral-induced inflammation (Figure 1C and D). After D-GalN/LPS injections, liver tissue injury was also measured (Figure 1C and D). However, no anti-HA antibody production was detected in these conditions, suggesting the absence of HA-specific adaptive immune response initiation (Figure 1E). Next, we asked whether local Th1 stimulation in the liver could initiate adaptive immune response against HA expressed by hepatocytes. Transient IL-12 expression in the liver of C57BL/6 mice has been reported as an inducer of chronic liver inflammation and injuries^31,32^. Thus, tamoxifen-treated RosaHAfl and HA/iCre mice received a hydrodynamic tail vein injection of a plasmid encoding murine IL-12 (pIL-12 HTVi) (Figure 1F). Histological analysis of liver tissue sections revealed pronounced lobular inflammation and penetrating portal infiltration after pIL-12 HTVi in both tamoxifen-treated RosaHAfl and HA/iCre mice, suggesting that the local inflammation induced was independent of HA expressed by hepatocytes (Figure 1G and H). As in Ad Ct i.v. and D-GalN/LPS conditions, pIL-12 HTVi treatment was not associated with anti-HA antibody production in mice expressing HA in the liver (Figure 1I). Altogether, these data showed that local non-specific liver injury and/or inflammation seem not to be sufficient to break the immune tolerance towards HA expressed by hepatocytes, and to give rise to HA-specific adaptive immune response.

**Figure 1.**
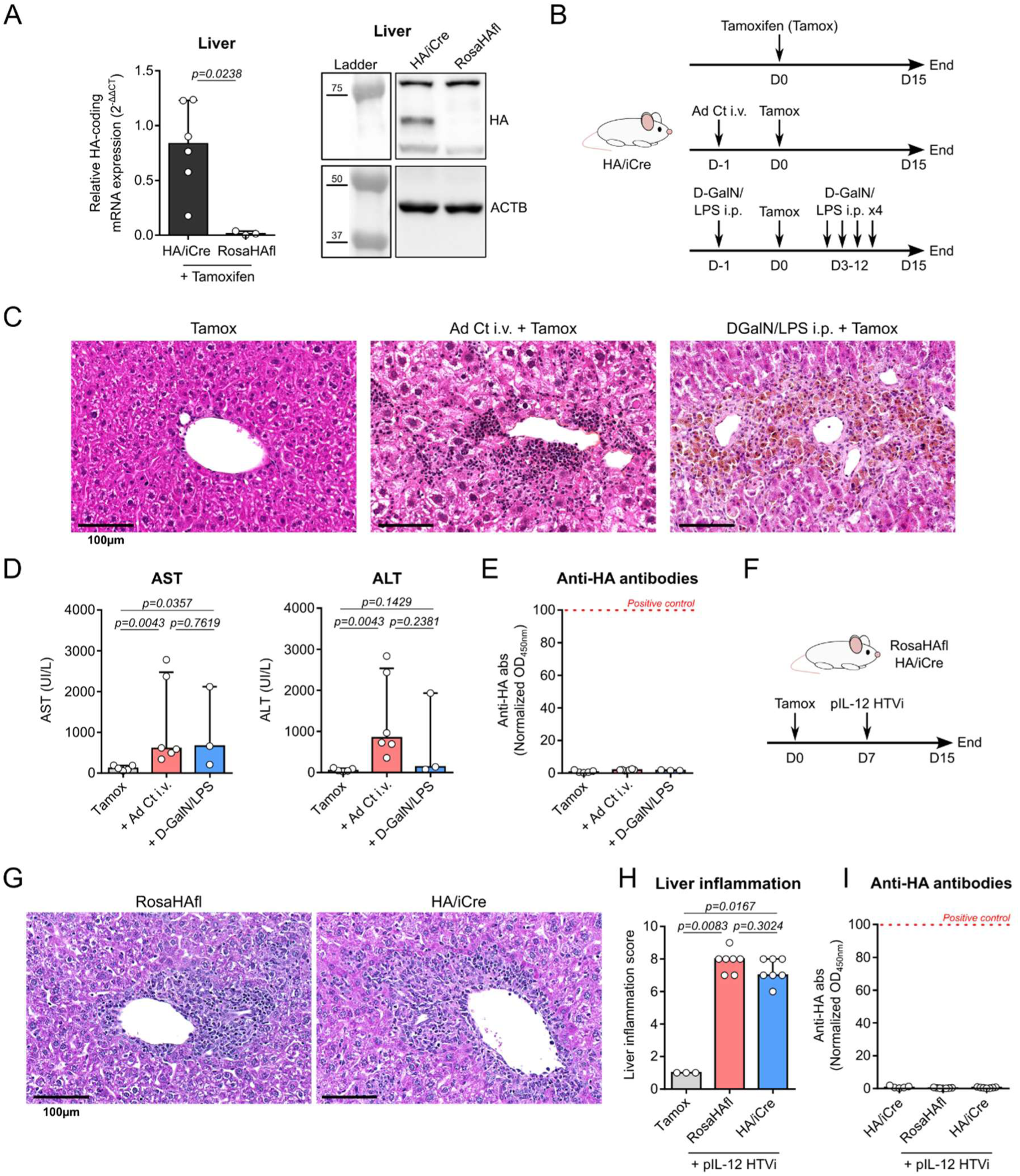
Addition of local inflammatory triggers in liver HA-expressing mice is not associated with anti-HA antibody production. **(A)** Relative HA expression in the liver of tamoxifen-treated HA/iCre (n=6) and RosaHAfl (n= 3) mice at the mRNA level by RT-qPCR (left) and at the protein level by western blot (right). ACTB was used as loading control. **(B)** Experimental design for the addition of liver inflammatory triggers. HA/iCre mice received either tamoxifen treatment alone (Tamox, n=5) or combined with intravenous (i.v.) injection of a control adenovirus (Ad Ct) one day before (n=5), or five intraperitoneal (i.p.) injections of D-GalN/LPS in a two week span. Mice were euthanized two weeks after the start of tamoxifen treatment. **(C)** Representative pictures of paraffin-embedded liver sections stained with HPS coloration from indicated conditions. **(D)** Transaminase (AST/ALT) levels in serum of mice. **(E)** Normalized anti-HA antibody rate in serum of mice. Positive control is indicated with a red dotted line. **(F)** Experimental design for hydrodynamic tail vein injection (HTVi) of IL-12-encoding plasmid (pIL-12) seven days after the start of tamoxifen on RosaHAfl (n=7) and HA/iCre (n=7) mice. Mice were euthanized one week after pIL-12 HTVi. **(G)** Representative pictures of paraffin-embedded liver sections stained with HPS coloration from indicated conditions. **(H)** Histological liver inflammation scoring analysis of liver tissue sections. **(I)** Normalized anti-HA antibody rate in serum of mice. Positive control is indicated with a red dotted line. Data are represented as median ± interquartile range in graphs **A**, **D**, **E**, **H** and **I**. Two-sided Mann-Whitney test was used for **A**, **D**, and **H**. p-values are indicated.

### Adenoviral-induced immunization against HA expressed in the liver is linked to a peripheral immune response

Previous AIH-like mouse models used intravenous injection of liver self-antigen-encoding adenovirus to generate a chronic autoreactive response in the liver with production of autoantibodies targeting hepatocellular antigens^15–18^. Despite the strong tropism of adenovirus for the liver, transduction of cells elsewhere could lead to the expression of the antigen outside the liver, making the origin of the primary antigen-specific response unclear in this context. Here, we analyzed the HA-specific immune response generated after i.v. injection of an adenovirus vector encoding the Cre recombinase (Ad Cre i.v.) in RosaHAfl or HA/iCre mice (Figure 2A). Ad Cre will induce HA expression in every transduced cell with the floxed Rosa26 HA sequence. For comparison, we analyzed the response in Ad Ct i.v. mice. We also tested RosaHAfl and HA/iCre mice after intramuscular immunization with Ad Cre or HA-encoding plasmid alone (HA i.m.) followed by tamoxifen treatment (HA i.m. + Tamox), which is known to induce HA-specific antibody production and HA-specific CD4 T cell accumulation in the liver of HA/iCre mice^3^ (Figure 2A). We detected HA mRNA expression in the liver of HA/iCre mice after intramuscular immunization and tamoxifen treatment, and after Ad Cre i.v. injection at similar levels, demonstrating liver transduction with the adenovirus (Figure 2B). Regarding HA-specific adaptive response, anti-HA antibody production was observed after intramuscular immunization against HA, as previously reported^3^, and with Ad Cre i.v. injection (Figure 2C). The i.v. injection of Ad Cre induced HA expression in the liver as well as anti-HA antibody production, whereas the i.v. injection of Ad Ct combined with tamoxifen-induced HA expression by hepatocytes was not sufficient to initiate HA-specific antibody production (Figure 1E). Both adenoviral transductions led to similar liver damages (Supplementary Figure 1). Thereby, we hypothesized that the HA-specific immune response is the consequence of HA expression by other cells in the periphery, outside the liver, after Ad Cre i.v. injection. This hypothesis is supported by a biodistribution study showing that, when injected intravenously, adenovirus vector rapidly accumulates in the liver then disseminates in the spleen^33^. We evaluated the relative HA or the relative Cre expression at the mRNA level by RT-qPCR in injected muscles of HA i.m. + Tamox mice or in the spleen of Ad Ct i.v. and Ad Cre i.v. mice (Figure 2D and E). As expected, intramuscular immunization was associated with HA expression in the injected muscles (Figure 2D). In the spleen, HA and Cre expression were detectable after Ad Cre i.v. injection suggesting that in addition to reaching the liver and transducing hepatocytes, Ad Cre also circulated in the periphery and induced HA expression in other cell types such as in the spleen (Figure 2E and supplementary Figure 1). These results suggest that B cell specific response towards a liver antigen is initiated through extrahepatic priming, even after hepatotropic adenoviral transduction.

**Figure 2.**
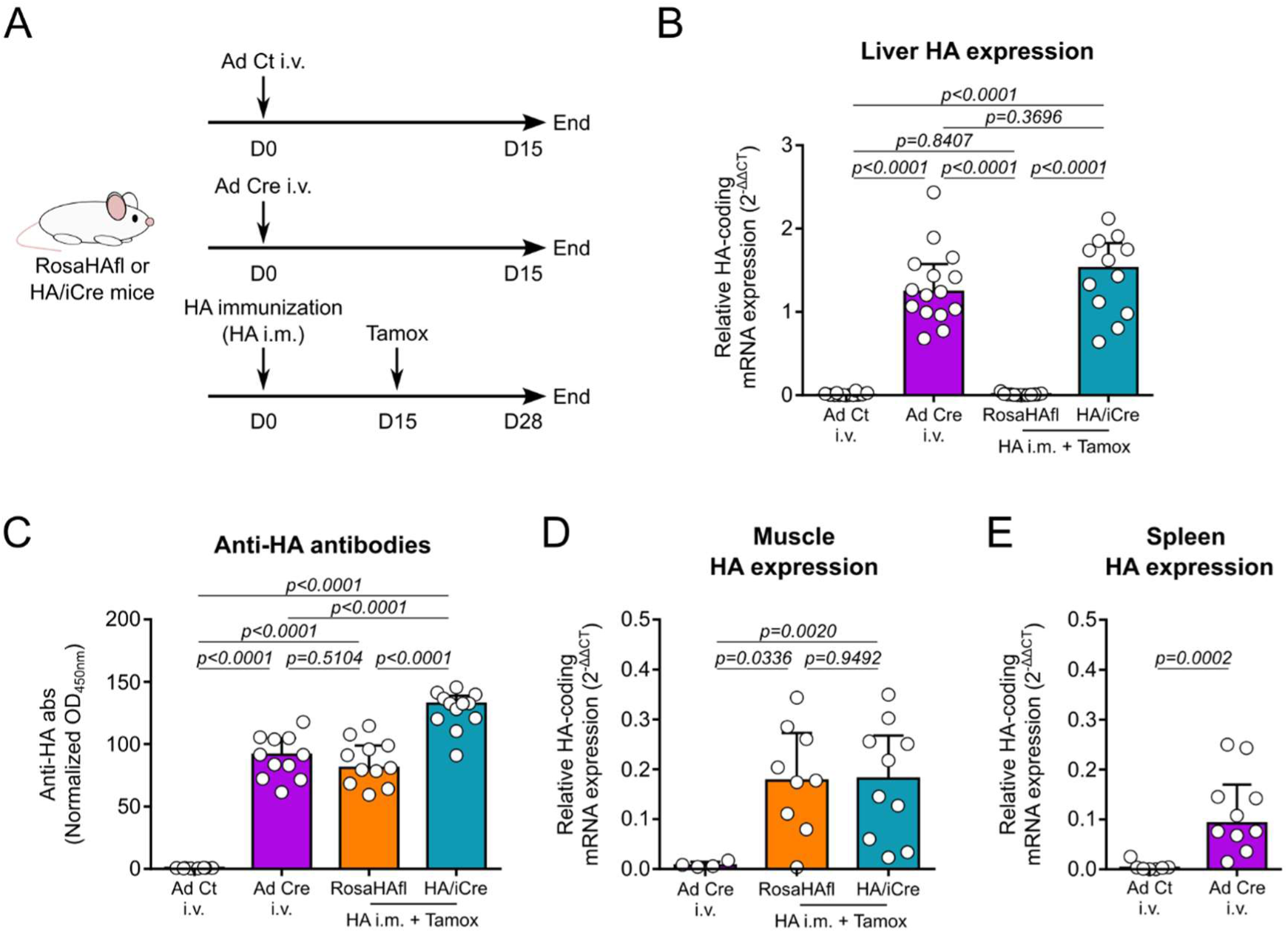
Intravenous injection of Ad Cre induces HA expression in the liver and in the periphery. **(A)** Experimental design for the induction of an immune response against HA. RosaHAfl or HA/iCre mice received either i.v. injection of a control adenovirus (Ad Ct, n=12), or a Cre-encoding adenovirus (Ad Cre, n=15), or intramuscular HA immunization followed by tamoxifen treatment (HA i.m. + Tamox, RosaHAfl, n=13; HA/iCre, n=12). Mice were euthanized two weeks after the last treatment. **(B)** Relative HA mRNA expression in the liver of mice. ACTB was used as loading control. **(C)** Normalized anti-HA antibody rate in serum of mice. **(D-E)** Relative HA mRNA expression in the tibialis anterior muscle **(D)** and in the spleen **(E)** of mice. ACTB was used as loading control. Data are represented as median ± interquartile range in graphs **B**, **C**, **D** and **E**. Two-sided Mann-Whitney test was used for **B**, **C**, **D** and **E**. p-values are indicated.

### Accumulation of antigen-specific T cells in the liver depends on local antigen expression after peripheral immunization

Since we generate an adaptive immune response against an antigen expressed in the liver, we wondered whether antigen-specific T cells are detectable in the liver tissue and what is their associated phenotype. We previously showed that HA-specific CD4 T cells could be tracked using PE-labeled I-A^d^ and I-E^d^ tetramers loaded with HA peptides, after a magnetic enrichment step^3^. Here, we used the same strategy to analyze HA-specific CD4 T cells in the spleen and the liver of Ad Ct i.v. and Ad Cre i.v. mice (Figure 3A and supplementary Figure 2). We also analyzed HA-specific CD8 T cells using PE-labeled H-2K^d^ tetramer loaded with HA peptide on splenocytes and liver non-parenchymal cells (NPCs) from Ad Ct i.v., Ad Cre i.v., and RosaHAfl or HA/iCre mice after immunization and tamoxifen treatment (HA i.m. + Tamox), which had not been achieved before (Figure 3A and supplementary Figure 2). We have previously described that HA-specific CD4 T cells were detected in the spleen after intramuscular HA immunization and were found in the liver only after tamoxifen-induced HA expression by hepatocytes, suggesting an active accumulation of antigen-specific CD4 T cells in the liver following local antigen expression^3^. Here, we detected HA-specific CD4 T cells in the spleen and the liver of mice after Ad Cre i.v. but not after Ad Ct i.v., suggesting similar process (Figure 3B). Regarding the CD8 T cell response, we were able to detect HA-specific CD8 T cells in the spleen of mice that were positive for anti-HA antibody production, which confirms complete adaptive immune response against HA in these models (Figure 3C, Figure 2C). In the spleen and the liver, no HA-specific CD8 T cells were detectable after tamoxifen treatment alone, as for HA-specific CD4 T cells^3^ (Supplementary Figure 3). In the liver, HA-specific CD8 T cells were detectable after Ad Cre i.v. injection but not after Ad Ct i.v. injection (Figure 3D). Marked accumulation of HA-specific CD8 T cells was found in the liver of HA immunized mice after local HA induction (HA/iCre, HA i.m. + Tamox, Figure 3D). Also, in the absence of hepatic HA expression, few HA-specific CD8 T cells were detectable in the liver upon immunization alone, which is different from HA-specific CD4 T cells^3^ (RosaHAfl, HA i.m. + Tamox, Figure 3D). While the frequency of HA-specific CD8 T cells per million liver NPCs seems not to be different after Ad Cre i.v. injection or intramuscular HA immunization alone (Figure 3D), Ad Cre i.v. mice display higher number of infiltrating HA-specific CD8 T cells along with higher number of total intrahepatic NPCs, CD4 T cells and CD8 T cells due to adenoviral transduction (Supplementary Figure 4). This suggests that, irrespective of liver HA expression, peripheral HA immunization leads to limited infiltration of HA-specific CD8 T cells in the liver. In addition, these results highlight the fact that accumulation of antigen-specific CD4 and CD8 T cells in the liver depends on local antigen expression after peripheral immunization.

**Figure 3.**
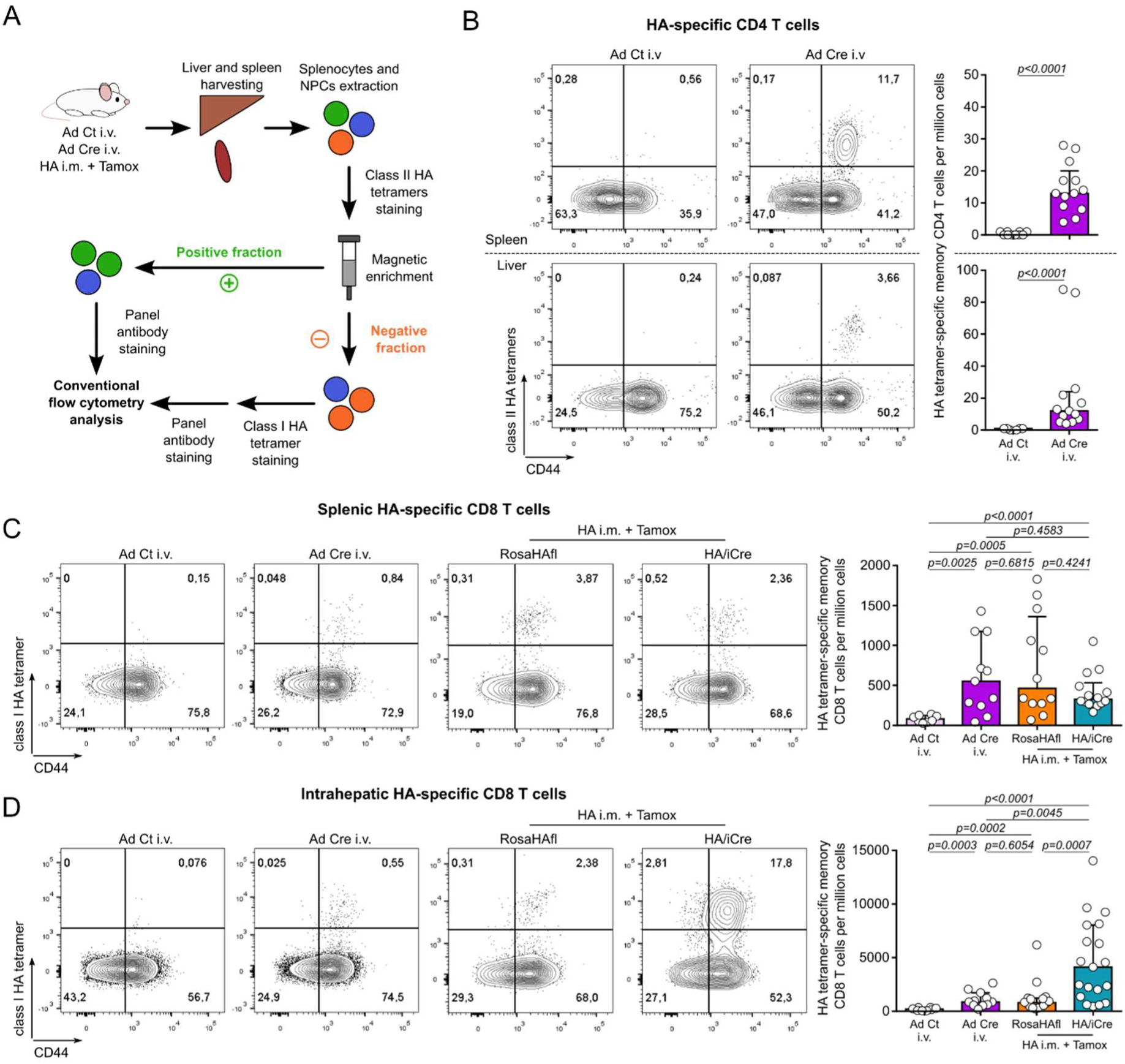
HA-specific CD4 and CD8 T cells accumulate in the liver following peripheral priming against HA and local HA induction. **(A)** Experimental design for tetramer staining on splenocytes and liver NPCs extracted from Ad Ct i.v. (n=10), Ad Cre i.v. (n=13), and RosaHAfl (n=16) or HA/iCre (n=19) mice that received HA i.m. + Tamox protocol. **(B)** Analysis of HA-specific CD4 T cells in the spleen (top) and in the liver (bottom). Contour plot representation of HA tetramer staining and CD44 expression in CD4 T cells from Ad Ct i.v. and Ad Cre i.v. mice (left). Frequency of HA-specific memory (CD44^high^) CD4 T cells per million cells (right). **(C-D)** Analysis of HA-specific CD8 T cells in the spleen **(C)** and in the liver **(D)** of Ad Ct i.v., Ad Cre i.v., and HA i.m. + Tamox (RosaHAfl or HA/iCre) mice. Contour plot representation of HA tetramer staining and CD44 expression in CD8 T cells (left). Frequency of HA-specific memory (CD44^+^) CD8 T cells per million cells (right). Data are represented as median ± interquartile range in graphs **B**, **C**, and **D**. Two-sided Mann-Whitney test was used for **B**, **C**, and **D**. p-values are indicated.

### PD-1 and TIGIT co-expression by T cells reflects local antigen-specific T cell response in the liver

Tetramer technology is highly specific but can be difficult and sometimes inapplicable for the study of antigen-specific T cells, like in AIH mouse models targeting human liver self-antigens CYP2D6 or FTCD. We therefore attempted to bypass the use of tetramers to detect a liver antigen-specific response by studying the phenotype of the reactive T cells in our models through conventional flow cytometry. Based on the previous work, we focused on the expression of PD-1 and TIGIT, found both in patients and in the mouse model at the transcriptomic level, and that seem to be specific of a chronic antigenic response in the tissue^3^. We first analyzed CD4 and CD8 T cells in the liver and in several lymphoid organs from RosaHAfl and HA/iCre mice that received intramuscular HA immunization followed by tamoxifen treatment, which represents a model that targets only the HA antigen without other exogenous antigenic stimulation. Among total intrahepatic CD4 and CD8 T cells, PD-1^+^ TIGIT^+^ populations were detected only in immunized mice expressing HA in the liver, more importantly for CD8 T cells (HA/iCre, Figure 4A and B). This phenotype was restricted to the liver, as PD-1^+^ TIGIT^+^ T cells were detected neither in the spleen, nor in the muscle-draining lymph nodes, or in the liver-draining lymph node (Figure 4B). While upon immunization alone a few HA-specific CD8 T cells were detected in the liver using class I HA tetramer, intrahepatic HA-specific CD8 T cells co-expressed PD-1 and TIGIT only when HA was locally induced (Figure 3D, Figure 4C). This highlights an association between PD- 1/TIGIT co-expression and local antigen reactivity as infiltrating HA-specific CD8 T cells are not expressing these markers until they encounter the antigen in the liver. Also, HA tetramer-specific and HA-reactive IFNγ-secreting CD4 and CD8 T cells were highly enriched in the intrahepatic PD-1^+^ TIGIT^+^ subsets (Figure 4D-F). Thus, HA-specific CD4 and CD8 T cells upregulate PD-1 and TIGIT by reacting against their antigen in the liver, which is consistent with the acquisition of an immuno- exhausted transcriptomic signature by HA-specific CD4 T cells following their recruitment in the tissue^3^.

**Figure 4.**
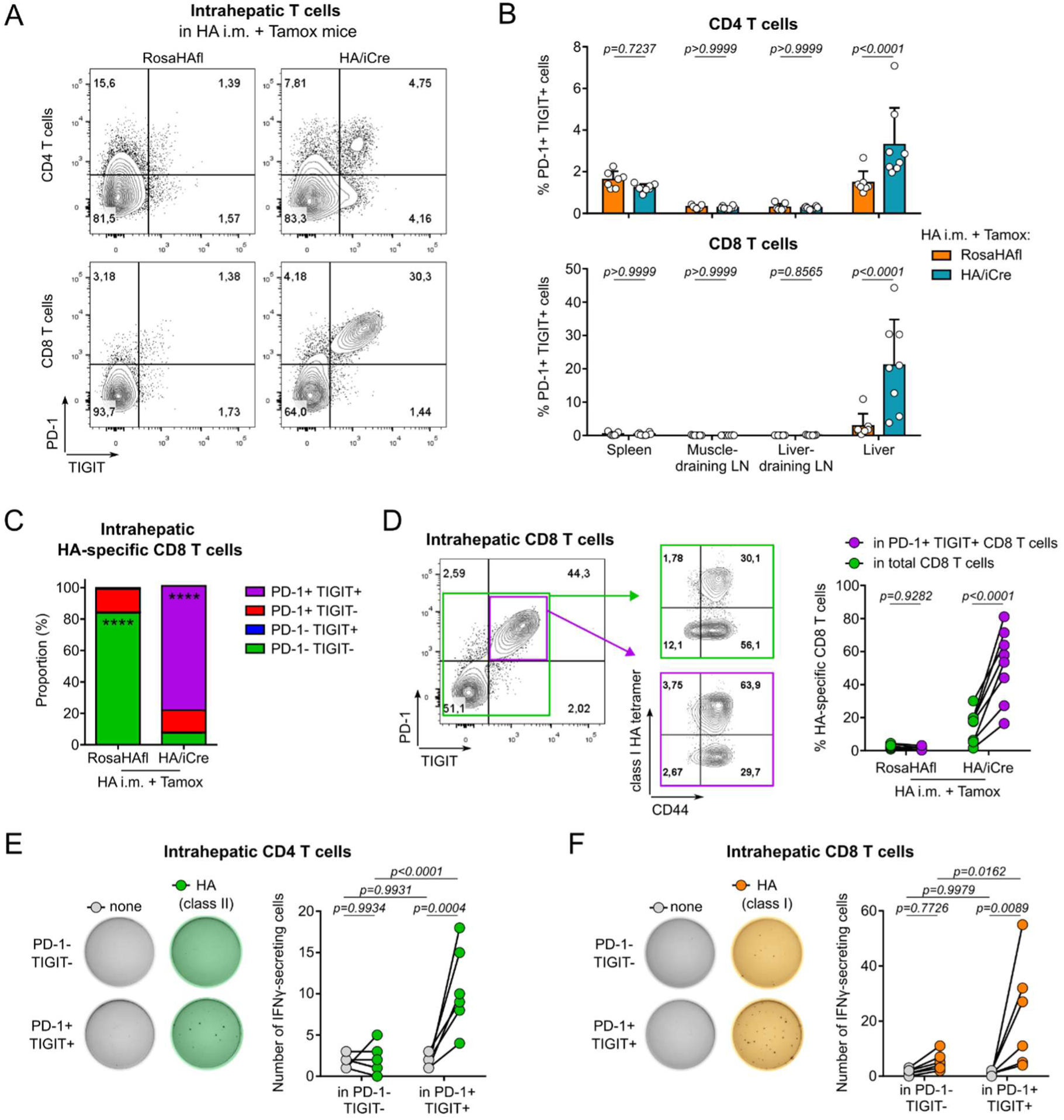
HA-specific T cell reactivity in the liver is marked with PD-1 and TIGIT co-expression by intrahepatic T cells. **(A)** Contour plot representation of PD-1 and TIGIT co-expression by CD4 T cells (top) and CD8 T cells (bottom) from the liver of RosaHAfl (left) and HA/iCre (right) mice following HA i.m. + Tamox protocol. **(B)** Analysis of PD-1/TIGIT co-expression by CD4 T cells (top) and CD8 T cells (bottom) from the spleen, muscle-draining lymph nodes (LN), liver-draining LN and liver of RosaHAfl (n=7) and HA/iCre (n=8) mice. **(C)** Proportion of PD-1/TIGIT expressing subsets among intrahepatic HA-specific CD8 T cells of RosaHAfl (n=7) and HA/iCre (n=8) mice. **(D)** Contour plot representation of HA-specific CD8 T cell frequency among total intrahepatic CD8 T cells (green) and PD-1^+^ TIGIT^+^ CD8 T cells (purple). Analysis of HA-specific CD8 T cell frequency between the two subsets in RosaHAfl and HA/iCre mice. **(E-F)** Representative pictures of HA-specific spot-forming assays with PD-1^-^ TIGIT^-^ or PD-1^+^ TIGIT^+^ CD4 T cells **(E)** and CD8 T cells **(F)** sorted from the liver of HA/iCre mice following HA i.m. + Tamox protocol (n=6, including one pool of two mice). Cells were co-incubated with splenocytes from a wild-type Balb/c mouse and were stimulated with CD4- stimulating HA peptides (class II) or with CD8-stimulating HA peptide (class I) or non-stimulated (none). Each black spot indicates one IFNγ-secreting cell (left). Number of IFNγ-secreting cells was compared between unstimulated and stimulated conditions (right). Data are represented as median in graph **C** and median ± interquartile range in graph **B**. Sidak’s multiple comparisons test was used for **B-F**. p-values are indicated. In **C**, ****: p<0.0001.

Based on the observation that PD-1^+^ TIGIT^+^ phenotype on T cells is associated with local antigenic reactivity, we expected to find accumulation of this phenotype on site where a local immune response is induced against the adenovirus-associated antigens. Thus, we investigated the PD-1/TIGIT co-expression by CD4 and CD8 T cells following intravenous injection of Ad Ct and Ad Cre. We observed that both Ad Ct and Ad Cre i.v. injections were associated with strong PD-1 and TIGIT co-expression by CD4 and CD8 T cells in the liver, showing local antigenic reactivity (Supplementary Figure 5A and 5B). PD-1^+^ TIGIT^+^ T cells were also found to a lesser extent in the spleen after Ad Ct and Ad Cre i.v. injection which was not the case after intramuscular immunization alone (Figure 4B and supplementary Figure 5B). This supports the fact that intravascular delivery of adenoviral vectors leads to transduction of hepatocytes as well as other cell types in the periphery^33^. Following Ad Cre i.v. injection, we found that both splenic and intrahepatic HA-specific T cells expressed PD-1 and/or TIGIT and that their proportion was enriched in total PD-1^+^ TIGIT^+^ subsets (Supplementary Figure 5C-G). Compared to HA/iCre mice in HA i.m. + Tamox protocol, HA-specific CD8 T cells represented only a little part of double positive T cells in the tissue, suggesting a marked concomitant T cell reactivity against adenovirus-associated antigens (Figure 4D and supplementary Figure 5E-G). This supports the idea that HA-specific response induced after Ad Cre i.v is linked to an extrahepatic priming against HA.

Altogether, these data demonstrate that T cell reactivity against an antigen in the tissue (liver or spleen) is associated with co-expression of PD-1 and TIGIT by T cells, providing a distinctive readout of local antigen-specific response in the liver.

### Intrahepatic tissue-activated CD4 and CD8 T cells are defined as PD-1^+^ TIGIT^+^ HLA-DR^+^ ICOS^+^ cells during AIH

We have previously detailed the immune signature of liver-derived autoreactive CD4 T cells found in the blood of autoimmune liver diseases patients by looking for intrahepatic TCR clonotypes into matched circulating memory CD4 T cells^3^. Yet, no high-dimensional phenotyping of AIH liver-infiltrating T cells has been performed. Here, we could have access to a liver sample from one AIH patient (referred as 10-004, See Methods) that underwent liver transplantation for severe disease. We performed immuno-phenotyping on tissue extracted lymphocytes using spectral flow cytometry. Among intrahepatic T cells, we could distinguish tissue-resident and non-resident T cells according to their CD69 expression^34–36^ (Figure 5A and supplementary Figure 6). Tissue-resident CD4 and CD8 T cells both showed strong co-expression of PD-1 and TIGIT compared to their CD69^-^ counterparts, suggesting local antigen reactivity in the tissue (Figure 5A). Dimension reduction analysis supported this result and revealed that tissue-activated CD4 and CD8 T cells are PD-1^+^ TIGIT^+^ HLA-DR^+^ ICOS^+^ T cells, a phenotype similar to circulating autoreactive CD4 T cells in AIH patients^3^ (Figure 5B-E). Also, these cells (notably CD8 T cells) express CD38, a marker that we have already described as upregulated by circulating T cells during AIH^2,3,37^, but also CD200 and/or CD161 which are associated to T peripheral helper and Th17 signatures in inflamed tissues, respectively^6,38,39^ (Figure 5D and E). These results were reproduced using unsupervised clustering analysis which distinguished PD-1^+^ TIGIT^+^ HLA-DR^+^ CD4 T cells into two subsets: a PD-1^hi^ ICOS^+^ CD25^-^ CD200^+^ subset (C03) and a PD-1^+^ ICOS^hi^ CD25^+^ CD200^-^ (C02, Supplementary Figure 7). These data represent the first high-dimensional phenotyping of liver-infiltrating T cells during AIH and support the idea that liver antigen reactivity is associated with PD-1 and TIGIT co-expression by intrahepatic CD4 and CD8 T cells.

**Figure 5.**
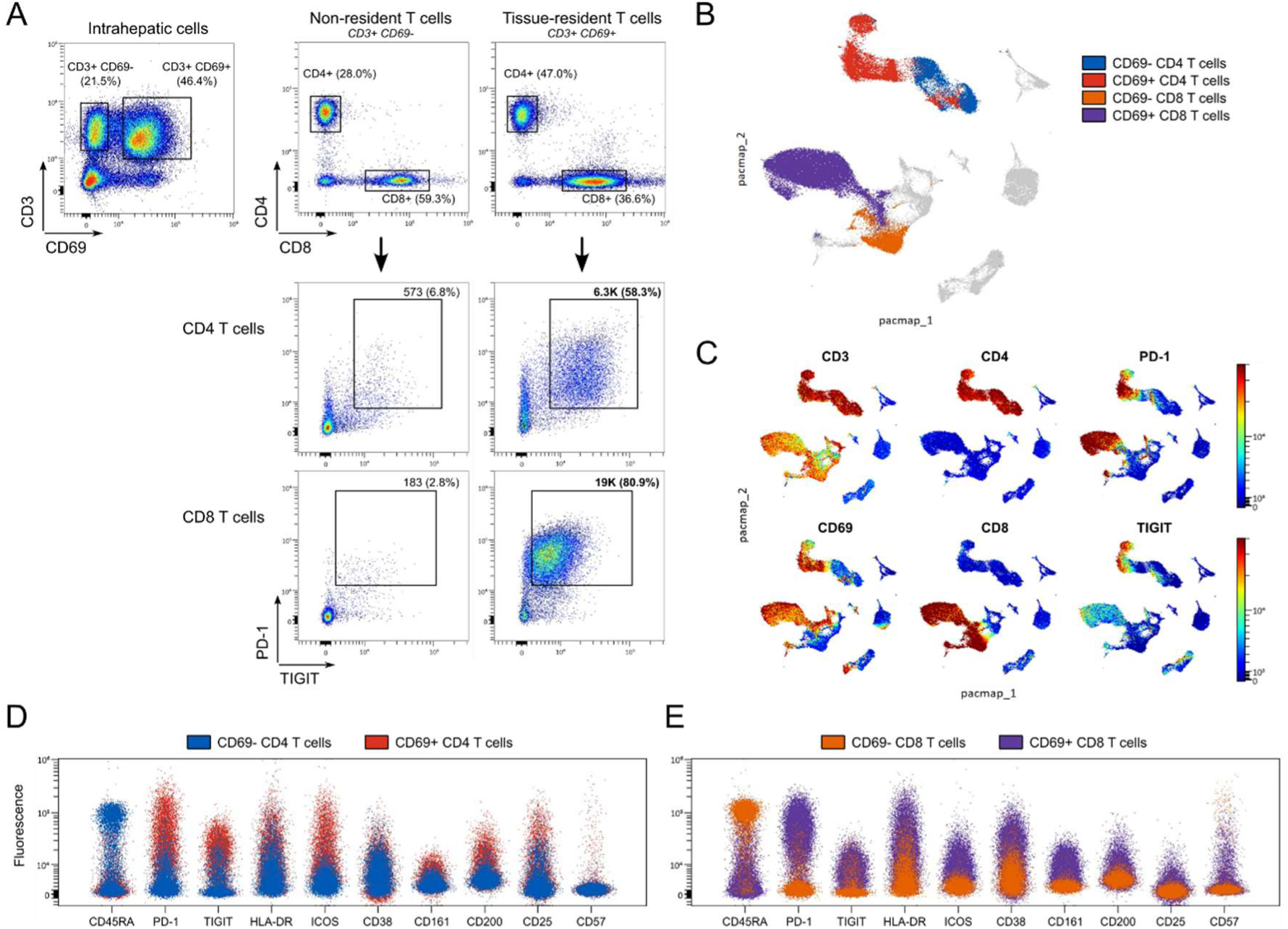
Immuno-phenotyping of intrahepatic T cells from AIH patient 10-004. **(A)** Pseudocolor dot plot representation of PD-1/TIGIT expression by CD4 and CD8 T cells from liver tissue-resident and non-resident T cells of AIH patient 10-004. **(B)** PaCMAP representation of intrahepatic lymphocytes, colored by CD4 and CD8 expression in CD3^+^ CD69^-^ and CD3^+^ CD69^+^ subsets shown in **A**. **(C)** CD3, CD69, CD4, CD8, PD-1 and TIGIT expression on PaCMAP representation. **(D)** Overlaid spectrum plot of CD69^-^ (blue) and CD69^+^ (red) CD4 T cells. **(E)** Overlaid spectrum plot of CD69^-^ (orange) and CD69^+^ (purple) CD8 T cells.

### Frequency of circulating PD-1^+^ TIGIT^+^ HLA-DR^+^ CD38^+^ CD8 T cells is associated with AIH disease activity

In AIH, the similarity between the phenotype of circulating autoreactive CD4 T cells and of intrahepatic CD4 T cells (from AIH patient 10-004) suggests that they share a common path. In the mouse model, local PD-1 and TIGIT co-expression by CD4 T cells following liver antigen induction supports the idea that circulating PD-1^+^ TIGIT^+^ CD4 T cells include liver-derived autoreactive CD4 T cells^3^. We have shown previously that circulating PD-1^+^ TIGIT^+^ HLA-DR^+^ CD4 T cells are more frequent in active AIH patients (AIHa) compared to patients in remission under treatment (AIHr) and non-alcoholic steatohepatitis (NASH) patients, suggesting that AIH disease activity could be associated with recirculation of these T cells^3^. To further support this hypothesis, we have now analyzed the phenotype of CD8 T cells in the blood of AIH patients. Using spectral flow cytometry data from the previously published study, we performed a new unsupervised clustering analysis on memory CD8 (mCD8) T cells from PBMCs of the same cohort of patients whose data were available (NASH, n=5; AIHa, n=10, AIHr, n=11)^3^ (Figure 6A, supplementary Figure 6 and supplementary Table 1). The analysis segregated mCD8 into 12 clusters (Figure 6B and 6C). Comparison of cluster frequencies revealed that three clusters (C7, C8, and C11 clusters) were significantly increased in AIHa patients compared to AIHr patients and to NASH patients (Figure 6D-F). Among these three clusters, the C7 was the most frequent and corresponded to PD-1^+^ TIGIT^+^ HLA-DR^+^ CD38^+^ ICOS^lo^ CD49d^+^ CD8 T cells, which was very similar to intrahepatic CD8 T cells from AIH patient 10-004 (Figure 5E and supplementary Figure 7) and circulating autoreactive CD4 T cells^3^ (Figure 6C, Figure 6G and supplementary Figure 8). The C11 cluster showed important proximity with the C7 cluster but was characterized by higher HLA-DR and ICOS expression, and CD25 expression (Figure 6C, Figure 6G and supplementary Figure 8). The C8 cluster differed from the two others with high expression of CD57, a marker of senescence^40^, that we have also found expressed by few liver activated CD8 T cells (Figure 5E and supplementary Figure 7) and circulating autoreactive CD4 T cells^3^ (Figure 6C, Figure 6G and supplementary Figure 8). Supervised analysis of HLA-DR, CD38 and ICOS expression in PD-1^+^ TIGIT^+^ mCD8 supported the idea that PD-1^+^ TIGIT^+^ HLA-DR^+^ and CD38^+^ mCD8 are enriched in active AIH patients (Supplementary Figure 9). We asked whether the frequency of these mCD8 subsets correlated with the liver injury of patients. As transaminases levels were used to classify AIH patients in active or in remission groups^41,42^ (Supplementary Table 1), we performed correlation analysis only with data from AIHa patients (Figure 6H). The C7 and C8 cluster frequencies significantly correlated with transaminase levels on this limited cohort of patients (Figure 6H). Also, supervised analysis revealed that, unlike PD- 1^+^ TIGIT^+^ HLA-DR^+^ mCD8, the frequency of circulating PD-1^+^ TIGIT^+^ CD38^+^ mCD8 significantly correlated with liver injury, which supports CD38 as a marker of AIH disease activity^37^ (Supplementary Figure 9). In comparison, C3, C5 and C10 cluster frequencies, which express PD-1 and TIGIT but lack HLA-DR, CD38 and ICOS expression, were not correlated with transaminase levels, supporting the idea that liver injury is associated with a specific PD-1^+^ TIGIT^+^ CD8 T cell subset (Figure 6C, Figure 6H and supplementary Figure 8). Therefore, PD-1^+^ TIGIT^+^ HLA-DR^+^ CD38^+^ memory CD8 T cells were found both in the liver and the blood during AIH, and their frequency in the blood of patients was associated with liver tissue injury. This suggests recirculation of tissue-activated intrahepatic T cells during the active phase of AIH.

**Figure 6.**
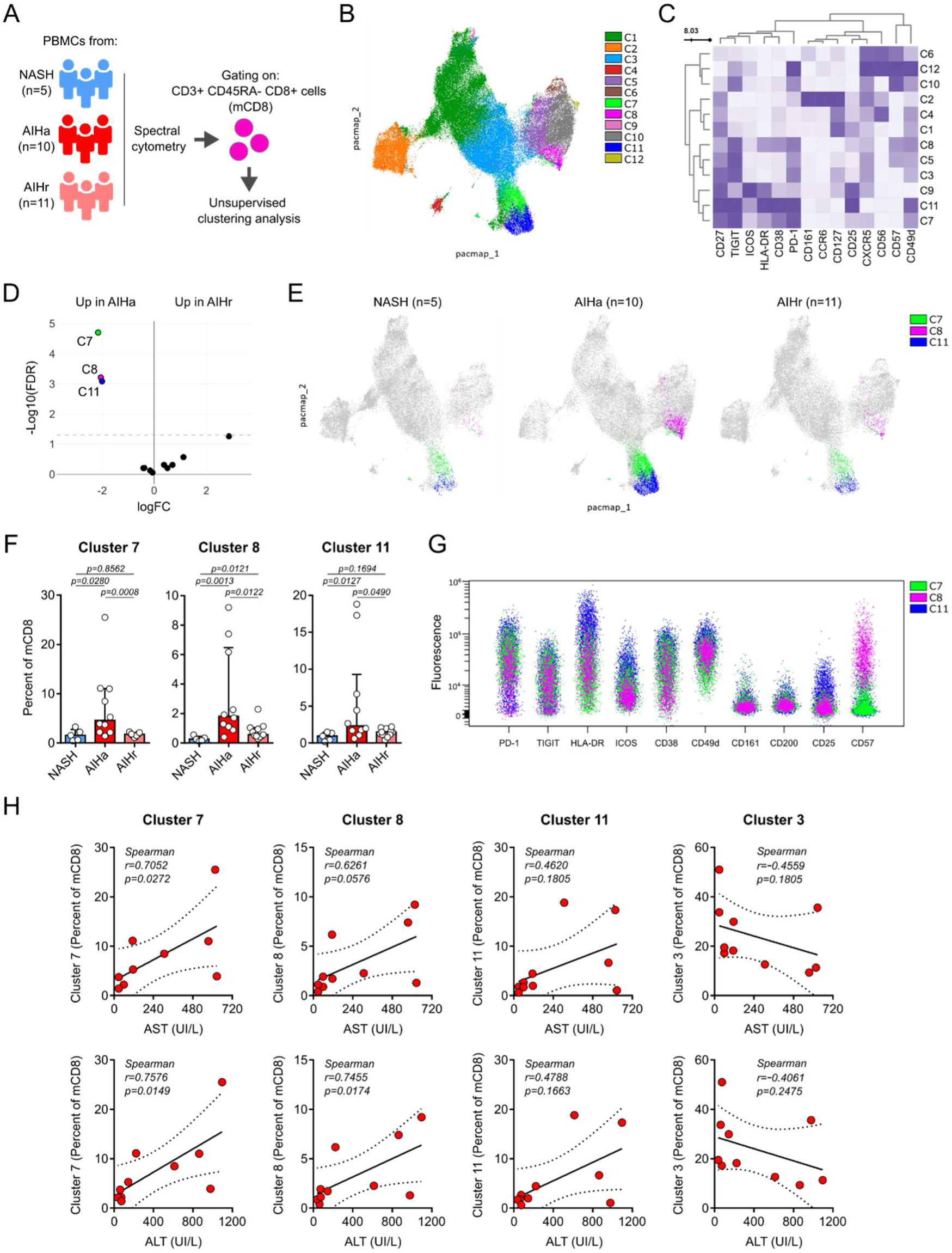
PD-1^+^ TIGIT^+^ HLA-DR^+^ CD38^+^ CD8 T cell subsets are enriched in the blood of active AIH patients. **(A)** Experimental design for immuno-phenotyping analysis of memory CD8 T cells from the blood of AIH patients in active phase (AIHa, n=10), in remission (AIHr, n=11) or from non-alcoholic steatohepatitis patients (NASH, n=5) as control. **(B)** PaCMAP representation of blood memory CD8 T cells colored by unsupervised FlowSOM clustering. **(C)** Surface marker heatmap of the clusters identified in **B**. **(D)** Differential cluster abundance between AIHa and AIHr patients. **(E)** PaCMAP representation shown in **B** colored by cluster 7 (C7, green), cluster 8 (C8, pink) and cluster 11 (C11, blue) in NASH (left), AIHa (middle) and AIHr (right) patients. **(F)** Analysis of C7, C8 and C11 frequencies in NASH, AIHa and AIHr patients. **(G)** Overlaid spectrum plot of C7, C8 and C11. **(H)** Correlation analysis of C7, C8, C11 and cluster 3 frequencies among blood memory CD8 T cells with AST (top) and ALT (bottom) rate in AIHa patients. Data are represented as median ± interquartile range in graph **F**. Two-sided Mann-Whitney test was used for **F**. Spearman correlation test was used for **H**. p- values are indicated.

## Discussion

In this study, using a non-TCR transgenic mouse model, we investigated the events required to initiate a complete adaptive immune response against an antigen (HA) expressed by hepatocytes, and we analyzed the phenotype of antigen-specific CD4 and CD8 T cells associated with this response. We demonstrated that production of HA-specific antibodies depends on the expression of HA outside the liver, as HA expression by hepatocytes alone or combined with local inflammatory triggers is not associated with specific humoral response. We showed that this peripheral immunization is required to find accumulation of HA-specific CD4 and CD8 T cells in the liver when HA is expressed locally. Using the mouse model and the liver sample from one AIH patient, we demonstrated that antigenic reactivity in the liver tissue is marked with PD-1 and TIGIT co-expression by intrahepatic CD4 and CD8 T cells. Deep immuno-phenotyping of circulating memory CD8 T cells from AIH and NASH patients revealed a PD-1^+^ TIGIT^+^ HLA-DR^+^ CD38^+^ CD8 T cell subset that shows high phenotypic proximity with tissue-activated T cells found in the AIH liver biopsy and that is enriched in the blood of active AIH patients and associated with liver injury. These findings revealed key interactions between the liver tissue and the periphery during T cell reactivity towards a liver antigen, from initiation steps to chronic active phase of the disease.

This study also provides an insightful mouse model to decipher the events involved in hepatocyte- antigen-specific immune response. Among the two models where we detected accumulation of HA-specific T cells in the liver, the protocol based on intramuscular immunization and tamoxifen-induced liver HA expression is a very simple model to target a single antigen expressed in the liver. We demonstrated that we could evaluate antigen expression and specific antibody production through easy assays, and that HA-specific CD4 and CD8 T cells could be detected using two distinct methods: class II and I HA tetramers, respectively, and *ex vivo* HA peptide stimulation. While these techniques are highly sensitive, the use of tetramers is still not available in the AIH-like mouse models targeting human liver self-antigens CYP2D6 or FTCD which limits the characterization of autoreactive T cells in these models. Then, as demonstrated in this work, PD-1 and TIGIT co-expression by T cells can be used as a reliable proxy to liver T cell reactivity for these models in the absence of other concomitant antigen reactivities (e.g. viral origin).

In this work, we found accumulation of antigen-specific T cells in the liver following either intramuscular immunization and tamoxifen-induced HA expression by hepatocytes, or intravenous injection of Ad Cre. As both conditions were associated with extrahepatic HA expression (in the muscle and the spleen respectively), this suggests that peripheral priming is essential for the accumulation of local hepatocyte-antigen-specific T cells. This hypothesis is supported by studies reporting that intravenous delivery of adenovirus vector leads to rapid Kupffer cells depletion and vector dissemination in the liver as well as in other organs^33,43^. Chronic AIH-like mouse model using human CYP2D6- or FTCD-encoding adenovirus vector may be initiated by extrahepatic transgene induction as well^15–18,23^. However, the response induced in these models appears to be highly influenced by genetic susceptibility and local inflammatory triggers, which suggests that in addition to potential extrahepatic priming, other events may regulate the initiation and evolution of an autoreactive response in the liver^16,23^. In our mouse model, further studies are needed to provide insights into long-term evolution of the liver antigen-specific immune response generated.

Once HA expression is induced in the liver of immunized mice, HA-specific CD4 and CD8 T cells accumulated locally in the tissue. In this condition, it is still unclear which T cell subset is the primary antigen recognizer. The fact that HA immunization alone (without liver HA expression) is associated with few HA-specific CD8 T cell-, and no HA-specific CD4 T cell-infiltration in the liver suggests that HA-specific CD8 T cells may be the first cells to detect HA expression at the surface of hepatocytes^44^. This is consistent with the fact that primed antigen-specific CD8 T cells take residency into several tissues, including the liver, where they mediate surveillance towards infection^45–48^. However, cross-presentation of hepatocyte antigen to specific CD8 T cells into the liver tissue or secondary lymphoid tissues has been reported^49,50^. Further investigation into the dynamics of antigen recognition in this model may inform on the primary response against a hepatocyte antigen.

The induction of HA in the liver of immunized mice was significantly associated with PD-1 and TIGIT co-expression by intrahepatic HA tetramer-specific T cells, which was also shared with tissue-resident T cells extracted from the liver biopsy of one AIH patient. This feature is consistent with the exhausted signature of liver self-antigen-specific CD4 T cells during chronic antigen exposure^3^. However, we can ask if this signature is restricted to the liver tissue. While liver antigen presentation has been associated with PD-1 and TIGIT expression by human and murine liver-infiltrating T cells during hepatocellular carcinoma and hepatitis B virus infection^9,10,14,44,51^, this feature is also shared with other tissues during autoimmunity and cancer^5,6,52–54^. Moreover, in this study, adenoviral transduction of cells in the spleen was also associated with local PD-1/TIGIT co-expression by CD4 and CD8 T cells independently of the hepatocyte antigen HA. Thus, PD-1 and TIGIT co-expression by T cells seems to reflect antigen reactivity in the tissue, including the liver.

In the blood of AIH patients, we have identified a PD-1^+^ TIGIT^+^ HLA-DR^+^ CD38^+^ memory CD8 T cell subset that is associated with disease activity and liver injury. This refines the CD8 T cell subset previously described as enriched in the blood of active AIH patients^2^. As discussed above, the analysis of PD-1^+^ TIGIT^+^ T cells must consider all possible tissue antigenic reactivities, indicating that these circulating PD-1^+^ TIGIT^+^ CD8 T cells may have reacted against their antigen in the liver, or elsewhere at some point. However, we showed that PD-1^+^ TIGIT^+^ CD8 T cell subsets that lack HLA-DR, CD38 and ICOS expression were not associated with liver injury, which suggests that an extended phenotype could be associated specifically with liver antigen reactivity. Furthermore, this CD8 T cell subset strongly mirrors the phenotype of circulating autoreactive CD4 T cells that represent dominant intrahepatic CD4 TCR clones^3^. Therefore, we can hypothesize that the circulating PD-1^+^ TIGIT^+^ HLA- DR^+^ CD38^+^ memory CD8 T cell subset could represent liver-derived T cells. It could reflect a high clonal expansion in the liver when the disease is active, and is consistent with a decrease of both the frequency of circulating PD-1^+^ TIGIT^+^ HLA-DR^+^ CD38^+^ CD8 T cells, and of the intrahepatic lymphocyte activity during remission^41^. Nevertheless, the reason for their presence in the blood of patients, just like autoreactive CD4 T cells, remains unclear. PD-1^+^ TIGIT^+^ HLA-DR^+^ CD38^+^ CD8 and autoreactive CD4 T cells could follow parallel trajectories from the liver during AIH. We can hypothesize that during active phase of the disease, T cell accumulation combined with liver damages in an already highly vascularized and fenestrated tissue could promote the escape of these T cells in the peripheral blood. Despite the unknown mechanisms mediating their presence the blood, recent studies have associated the frequency of similar circulating T cells with disease onset during rheumatoid arthritis^55^, and with treatment response during AIH^37^. Therefore, our findings could provide new immune-based markers to monitor AIH activity and enable a deeper characterization of liver-activated CD4 and CD8 T cells isolated from the blood of patients.

## Methods

### Patients

All the patients eligible signed a written informed consent prior to inclusion into a bio-bank of samples of AILD patients (BIO-MAI-FOIE) maintained in Nantes University Hospital which obtained regulatory clearance from the biomedical research Ethics Committee (COMITE DE PROTECTION DES PERSONNES OUEST IV-NANTES CPP) and all required French Research Ministries authorizations to collect biological samples (Ministère de la Recherche, ref MESR DC-2017-2987). The biobank is supported by the HEPATIMGO network promoted since 2017 (RC17_0228) by Nantes University Hospital and is a prospective multi-centric collection managed by the Biological Resource center of the CHU of Nantes. All data collected were treated confidentially and participants are free to withdraw from the study at any time, without explanation and without prejudice to their future care. It was granted authorization from the CNIL: 2001209v0. All AIH patients included in this study had a simplified diagnostic score superior or equal to 6 according to the simplified scoring system for AIH of the international autoimmune hepatitis group (IAHG)^42,56^. Active AIH (AIHa) patients are untreated patients with new onset AIH patients enrolled at diagnosis prior any treatment initiation as previously described^41^, and AIH patients under standard treatment but who do not normalize the transaminases (AST and ALT) and/or the serum IgG levels or are under relapsing event (Supplementary Table 1). AIH patient 10-004 was a 17 years old female who has undergone liver transplantation for severe AIH disease (IgG: 52,1g/L, AST: 305,0IU/L, ALT: 222,0UI/L, AIH score: 7). Remission AIH (AIHr) patients are defined biochemically by a normalization of the transaminases and the IgG levels, according to the most recent European clinical practice guidelines on AIH^42^. All NASH (Non-Alcoholic SteatoHepatitis) patients had histological evidence of NASH and had dysmetabolic syndromes (Supplementary Table 1). No sex analysis was carried out, as AILD are rare diseases that predominantly affects women^42,57^. This study was carried out in accordance with the Principles of International Conference on Harmonisation (ICH) Good Clinical Practice (GCP) (as adopted in France) which builds upon the ethical codes contained in the current version of the Declaration of Helsinki, the rules and recommendations of good international (ICH) and French clinical practice (good clinical practice guidelines for biomedical research on medicinal products for human use) and the European regulations and/or national legislation and regulations on clinical trials.

### Cell extraction and spectral flow cytometry analysis

Blood samples from patients were collected and PBMCs were isolated through Ficoll density gradient centrifugation. Liver sample was obtained after AIH patient 10-004 undergone liver transplantation for severe AIH. Liver sample was cut into small pieces and digested with collagenase IV (C5138, Sigma) at 37°C for 15min. Then, liver samples were crushed using gentleMACS Dissociator (Miltenyi) followed by manual grinding on 100µm filter. Intrahepatic mononuclear cells were isolated through Ficoll density gradient centrifugation.

For surface staining, 1x10^6^ PBMCs or intrahepatic mononuclear cells were incubated 20min with a mix of antibodies and then washed prior analysis on Cytek Aurora. All antibodies used are described in the supplementary Table 2. Flow cytometry data from patients were analyzed using Omiq software (Dotmatics, USA). CCR2, LAG-3, CXCR3 and BTLA markers were excluded from analysis as their associated fluorescence were not interpretable. Concerning analysis of blood memory CD8 T cells, subsampling of viable memory CD8 T cells (CD3^+^ CD4^-^ CD8^+^ CD45RA^-^ cells, Supplementary Figure 6) was performed (7x10^3^ per patients). Clustering was performed using FlowSOM to generate 12 clusters. Dimension reduction was represented using PaCMAP. edgeR was used to identify significant differential cluster abundance between groups of patients. Concerning analysis of intrahepatic T cells, 100x10^3^ lymphocytes (FSC/SSC) were subsampled and analyzed through supervised or unsupervised strategy. Intrahepatic resident T cells (CD3^+^) were differentiated from non-resident T cells according to their CD69 expression and then separated into CD4 and CD8 T cells (Supplementary Figure 6). Clustering was performed using FlowSOM to generate 20 clusters from total intrahepatic lymphocytes. Dimension reduction was represented using PaCMAP.

### Mice and procedures

All procedures including mice were performed in accordance with guidelines and regulations approved by the regional ethical committee for animal care and use and by the Ministère de l’enseignement supérieur et de la recherche (agreements APAFIS #2054, #28582 and #43529). Mice were housed at the UTE IRS-UN animal facilities (Nantes, FRANCE) in specific pathogen-free conditions, at a maximum of five mice per individually ventilated cage in 12h:12h light/dark cycle and under standard conditions of temperature (21-24°C) and humidity (40-60%). Mice were fed *ad libitum* with continuous access to tap water, and each cage was composed of environmental enrichment material within the limits of available stocks. As described before^3^, homozygous *Rosa26^tm(HA)1Libl^* (RosaHAfl) mice were kindly provided by R. Liblau (Toulouse, FRANCE), and heterozygous transthyretin (TTR)-inducible Cre (*TTR-Cre-ind*) mice were back-crossed on a Balb/c background for at least 10 generations (TAAM, CDTA CNRS Orléans, FRANCE)^27,28^. Cross-breeding of RosaHAfl and *TTR-Cre-ind* mice resulted in *Rosa26^tm(HA)1Libl^/TTR-Cre-ind* (HA/iCre). At 3 weeks old, mice were genotyped for Cre-coding gene thanks to small tail biopsies using Cre primers (Supplementary Table 3) Each experiment used male and female eight to twelve-weeks-old mice with co-housing of experimental and control animals. At the end of experiments, mice were anaesthetized and euthanized by cervical dislocation.

Tamoxifen treatment was performed by feeding mice with tamoxifen dry food (0.5g/kg tamoxifen + 5% saccharose; Safe, FRANCE) for 14 days in free access or with five intraperitonial injections (i.p) of 1mg of tamoxifen (T5648, Sigma) diluted in corn oil (C8267, Sigma) in two weeks span. To induce liver inflammatory triggers, mice received either intraperitoneal co-injections of D(+)-galactosamine hydrolochloride (D-GalN, 400mg/kg, G0500, Sigma) and lipopolysaccharide (LPS, 2µg/kg, L2880, Sigma) five times in two weeks span, or hydrodynamic tail vein injection (HTVi) of a murine IL-12- coding plasmid (#123139, Addgene) (5-20µg diluted in 9% body weight (mL) of PBS)^58–60^. Delivery of adenoviral (Ad) vectors was performed through intravenous injection (i.v.) of 3,0.10^9^ infectious particle of Ad Ct (Ad CMV rAFP, Ad CMV GFP or Ad CMV hAFP, produced by INSERM UMR 1089 CPV facility, Nantes, FRANCE) or Ad Cre (Ad CAG Cre, produced by INSERM UMR 1089 CPV facility, Nantes, FRANCE).

Intramuscular immunization against HA was performed with two different materials: one intramuscular (i.m.) injection of 1,5.10^9^ infectious particle of Ad Cre (diluted in 10-20µL PBS) in *tibialis anterior* of one posterior leg or i.m. injection of 50µg of plasmid CMV HA vector (diluted in 30µL PBS) in *tibialis anterior* of both posterior legs, twice three weeks apart. As described before^3^, plasmid CMV HA vector, HA sequence (Influenza A virus (A/Puerto Rico/8/1934(H1N1), Gene ID: 956529)) was synthesized (GenScript, NETHERLANDS) and cloned in pCMVβ vector (Clontech Laboratories) in place of β-galactosidase gene.

### RNA extraction, reverse transcription, and quantitative PCR

Liver, spleen and muscle sample were collected and freeze at -80°C. Total RNA was extracted from organ tissue using TRIzol reagent (15596026, ThermoFisher Scientific) and purified with RNeasy Mini Kit (74106, Qiagen) according to the manufacturer protocol. Reverse transcription was performed using 2µg of total RNA incubated at 70°C for 10min with poly-dT24 20µg/mL (Eurofins Genomics), 8mM DTT (18057018, ThermoFisher Scientific) and 20mM of each dNTP (10297018, ThermoFisher Scientific). After a brief 5min incubation at 4°C, first strand buffer 5X (Y00146, Invitrogen), 200U of M-MLV reverse transcriptase (18057018, ThermoFisher Scientific) and 40U of RNAse OUT inhibitor (10777019, Invitrogen) were added and incubated at 37°C for 1h followed by 15min at 70°C. Real-time quantitative PCR was performed using the ViiA 7 Real-Time PCR System and Power SYBR Green PCR Master Mix (4368708, ThermoFisher Scientific). Primers used for HA, Cre and ACTB relative mRNA expression analysis are listed in the supplementary Table 3. Relative HA mRNA expression was determined with 2^-ΔΔCT^ calculation, taking ACTB as reference gene and liver cDNA from one conserved Ad Cre i.v. mouse as positive control. Relative Cre mRNA expression was determined with 2^-ΔCT^ calculation, taking ACTB as reference gene.

### Western blot

Total proteins were extracted from liver samples via RIPA buffer treatment. 25µg of protein were denatured at 95°C for 5min in Laemmli Sample Buffer (161747, Bio-Rad) with DTT 0.1M (10197777001, Sigma). Preparation was separated by SDS-PAGE on Mini-PROTEAN TGX Precast Protein Gels (4561036, Bio-Rad) in migration buffer (Tris 15g/L; Glycine 72g/L; SDS 10g/L) and transferred onto a PVDF membrane with the Trans-Blot Turbo Transfer System (Bio-Rad). The membrane was stained with Ponceau S solution (P7170, Sigma) and was cut horizontally above the 50kDa molecular weight ladder. Both parts of the membrane were blocked for 2h with a blocking solution (TBS; Tween 20 0.1%; Skim milk 5%), followed by overnight incubation at 4°C with anti-HA antibody (rabbit polyclonal antibody; 11684-T62, Sinobiological) for the upper membrane and anti-mouse ACTB antibody (mouse monoclonal antibody; 3700, Cell Signaling) for the lower membrane, as primary antibodies. The membrane was washed and incubated 1h at room temperature with peroxidase-conjugated donkey anti-rabbit IgG (H+L) antibody (E-AB-1080-120, Clinisciences) and peroxidase-conjugated goat anti-mouse IgG+IgM (H+L) antibody (115-036-068, Jackson ImmunoResearch). Revelation was performed by Electrochemioluminescence Super Signal West Pico (34577, ThermoFisher Scientific) according to the manufacturer instructions. Imaging and analysis of western blots were performed on the ChemiDoc MP Imaging System (Bio-Rad). Total protein extract from liver sample of one conserved Ad Cre i.v mouse and one wild-type mouse was used as positive and negative controls respectively.

### Histological analysis and serum transaminase dosage

Livers were sampled and lobes were fixed in formol 4% (11699404, VWR) for 24h at room temperature. Dehydration in differential absolute ethanol baths, embedding in paraffin and staining of paraffin-embedded sections (3µm) with hematoxylin-phloxin-saffron (HPS) coloration were performed by the IBISA MicroPICell facility (Biogenouest; Nantes, FRANCE). Slides were scanned and observed using NanoZoomer S360 (Hamamatsu) and NDP Scan software. As previously described^3^, liver inflammation scoring analysis was performed blindly and based on established hepatitis grading^61^ simplified to portal inflammation score (0 to 4), lobular inflammation score (0 to 4) and presence of interface hepatitis (0 to 1), giving a minimum score of 0 and maximum score of 9.

At the end of each experiment, blood was sampled. Serum was harvested after centrifugation step and stored at -80°C. Dosage of aspartate aminotransferase (AST) and alanine aminotransferase (ALT) in the serum of mice was performed by the Centre Hospitalier Universitaire of Nantes (FRANCE).

### Anti-HA antibody detection

Anti-HA antibody production was tested through ELISA as previously described^3^. Briefly, plates were coated with HA protein (1µg/mL diluted in coating buffer (Na_2_CO_3_ 0.05M; NaHCO_3_ 0.05M; pH 9.2), 11684-V08H, SinoBiological) and incubated at 4°C overnight. After washing steps, plates were saturated with PBS 1X Tween 20 0.05% BSA 1% for 2h at 37°C. Plates were washed and serum samples were added to wells in duplicate for three dilutions: 500, 1000 and 5000. After an incubation of 2h at 37°C, plates were washed and detection antibody was added (0,4ng/mL, peroxidase-conjugated goat anti-mouse IgG+IgM (H+L) polyclonal antibody, 115-036-068, Jackson ImmunoResearch) and incubated for 1h at 37°C. Revelation step was performed with TMB Substrate Reagent (555214, BD Biosciences) and stopped at 1min30sec by addition of H_2_SO_4_ 0.5M. Optical densities (OD) were determined using a Spark 10M Infinite M200 Pro plate reader (TECAN). Analysis was performed by blank reduction, mean calculation and normalization of OD values to OD value of a conserved positive control serum (Ad Cre i.v. mouse), for each corresponding dilution. Mean of normalized OD values for each dilution was calculated and compared between samples and conditions.

### Cell preparation

Spleens were sampled and splenocytes were extracted by mechanical dissociation on 40µm filter in red blood cell lysis buffer (NH_4_Cl 155mM; KHCO_3_ 10mM; EDTA 1mM; Tris 17mM per 1L of sterilized water). Muscle-draining lymph nodes and liver-draining lymph nodes were sampled and cells were extracted by mechanical dissociation on 40µm filter. Livers were perfused with HBSS 1X and non-parenchymal cells (NPCs) were isolated as previously described^3,62^. Briefly, livers were cut and digested with collagenase IV (C5138, Sigma) at 37°C for 15min, and then crushed on 100µm filter. After centrifugation steps, liver NPCs were enriched by 40%:80% Percoll (GE17-0891-01, Sigma) density gradient centrifugation and red blood cells lysis step.

### Tetramer enrichment, antibody staining and conventional flow cytometry

10 to 20.10^6^ splenocytes and 3.10^6^ liver NPCs were stained with both I-A^d^-HA peptide (HNTNGVTAACSHE) and I-E^d^-HA peptide (SFERFEIFPKE) PE-labelled tetramers (NIH Tetramer Core Facility; Atlanta, USA) at room temperature during 1h. Cells were washed and stained with magnetic anti-PE microbeads (130-048-801, Miltenyi) at 4°C during 15min. Cells were washed and tetramer-specific T cells were enriched using magnetic MS columns (130-042-201, Miltenyi). Positive fraction was kept for HA-specific CD4 T cells analysis. Negative fraction was stained with H-2K^d^-HA peptide (IYSTVASSL) PE-labelled tetramer (NIH Tetramer Core Facility; Atlanta, USA) at room temperature during 1h for HA-specific CD8 T cells analysis. All cells were washed and viability staining (LIVE/DEAD Fixable Aqua Dead Cell Stain kit, dilution 1:1000, L34957, ThermoFisher Scientific) was performed at 4°C during 15min, protected from light. Cells were washed and extracellular staining was performed at 4°C during 20min protected from light, using fluorochrome-coupled anti-mouse antibodies (Supplementary Table 2). Fluorescence was measured on BD FACSCantoII or BD FACSAriaII (BD Biosciences; Mountain View, USA). HA-specific memory CD4 and CD8 T lymphocytes were defined as: LIVE/DEAD^-^ CD19^-^ CD4^+^ (or CD8^+^) CD44^high^ tetramer^+^ cells, as described in supplementary Figure 2. FlowJo software (TreeStar Inc) was used to analyze flow cytometry data.

### HA-specific IFNγ-secretion assay

To detect HA-reactive cells ELISpot assay was performed following manufacturer BD™ ELISPOT Set Instruction Manual. Plates (S2EM004M99, Merck Millipore) were coated with 5µg/mL of BD NA/LE Purified anti-mouse IFN-γ capture antibody (551881, BD Biosciences) and incubated overnight at 4°C in sterile conditions. Then, wells were blocked with RPMI medium (supplemented with 10% FBS 1% Penicillin/Streptomycin, 1% L-Glutamine) for 2 hours at room temperature. After washing step, stimuli and cells were added and plates were incubated at 37°C overnight in 5% CO2 humidified incubator. Biotinylated anti-mouse IFN-γ detection antibody (2µg/mL, 551881, BD Biosciences) was added and incubated for 2 hours at room temperature and followed by incubation with HRP Streptavidin (557630, BD Biosciences) for 1 hour at room temperature. Finally, revelation step was performed following AEC Substrate Set protocol (551951, BD Biosciences) and reaction was stopped 30 minutes after with deionized water and wells were washed several times. Plates were air-dried at room temperature and were red on IRIS 2 reader (Mabtech). In this work, we sorted PD-1/TIGIT double positive and double negative CD4 and CD8 T cells from the liver of tamoxifen-treated immunized HA/iCre mice to test their reactivity against HA. For each well, 2,0.10^3^ sorted CD4 subset or 3,5.10^3^ sorted CD8 subset were added to 200.10^3^ feeder splenocytes from one wild-type mouse. HA stimulation was performed by adding 5µg/mL HNTNGVTAACSHE and 5µg/mL SFERFEIFPKE peptides (Sigma) to stimulate HA-specific CD4 T cells or 5µg/mL IYSTVASSL peptide (Synpeptide) to stimulate HA-specific CD8 T cells. Culture medium was used as negative control and wild-type splenocytes alone were incubated with Dynabeads CD3/CD28 stimulator (11456D, Gibco) for technical positive control.

### Statistical analysis

Statistical comparisons were performed using GraphPad Prism software V.6 (GraphPad Software, La Jolla, CA, USA). p-value <0.05 after adjustment were considered significant.

## Acknowledgements

We thank the biological resource center for biobanking (CHU Nantes, Hôtel Dieu, Center de ressources biologiques (CRB), Nantes, F-44093, France (BRIF: BB-0033-00040)). We thank all the members of the HEPATIMGO network. We thank Dr. Sarah HABES, Dr. Maëva SALIMON, and Dr. Annie LIM for the inclusion of new AIH patients. We thank the UTE IRS-UN animal facility of the SFR Santé François Bonamy (Nantes Université, INSERM UAR016, CNRS UAR3556). We acknowledge the MicroPICell core facility (SFR Bonamy, BioCore, Inserm UMS 016, CNRS UAR 3556, Nantes, France), member of the Scientific Interest Group (GIS) Biogenouest, IBISA, and the national infrastructure France-Bioimaging supported by the French national research agency (ANR-10-INBS-04). We thank TaRGeT laboratory, the GTI core and ViVeM center (Nantes Université, CHU Nantes, INSERM, TaRGeT, F- 44000 Nantes, France) for viral vector production and help in ELISpot plate reading. We thank the NIH tetramer core facility for provision of MHC-peptide complexes. Supported by the Agence Nationale de la Recherche (ANR-19 CE17-0024, ANR-24 CE15-7123), the patient association Association pour la Lutte contre les maladies inflammatoires du foie et des voies biliaires (ALBI), the Fondation Maladies Rares, the Région pays de la Loire, the LabEx IGO program (n° ANR-11-LABX-0016) funded by the Investment into the Future French Government program managed by the Agence Nationale de la Recherche (ANR). This work was supported by institutional grants from INSERM.

## Author Contributions

T. G. performed the experiments, analyzed the data and wrote the manuscript; A. C. performed the experiments and analyzed the data; J-P. J. performed the experiments; P-J. G. performed the experiment and analyzed the data; F. R. and V. H. performed the experiments; C. C. managed human AIH samples collection and patient data base; M. K., E. B-J., L. E., A. L., C. S., M. S., F. T., S. L. and E. D. M. provided human AIH samples and critical insight in AIH pathology; F. V. managed human AIH samples collection; A. N. provided the RosaHAfl mouse strain; S. B. provided critical insight in the study design; J-F. M. provided critical insight in histology analysis; J. G. provided human AIH samples, critical insight in AIH pathology and direction in the study design; S. C. designed the study, supervised data analysis and wrote the manuscript; A. R. designed the study, supervised data analysis, performed the experiments and wrote the manuscript. All authors reviewed and approved the manuscript.

## Data Availability

Raw numbers for charts and graphs are available in the XXXX file provided with this paper. The FCS data from human patients’ samples are available in the following repository: https://xxxxxxx/. The FCS data from murine experiments are available from the corresponding author upon reasonable request.

---

**Supplementary Figure 1.**
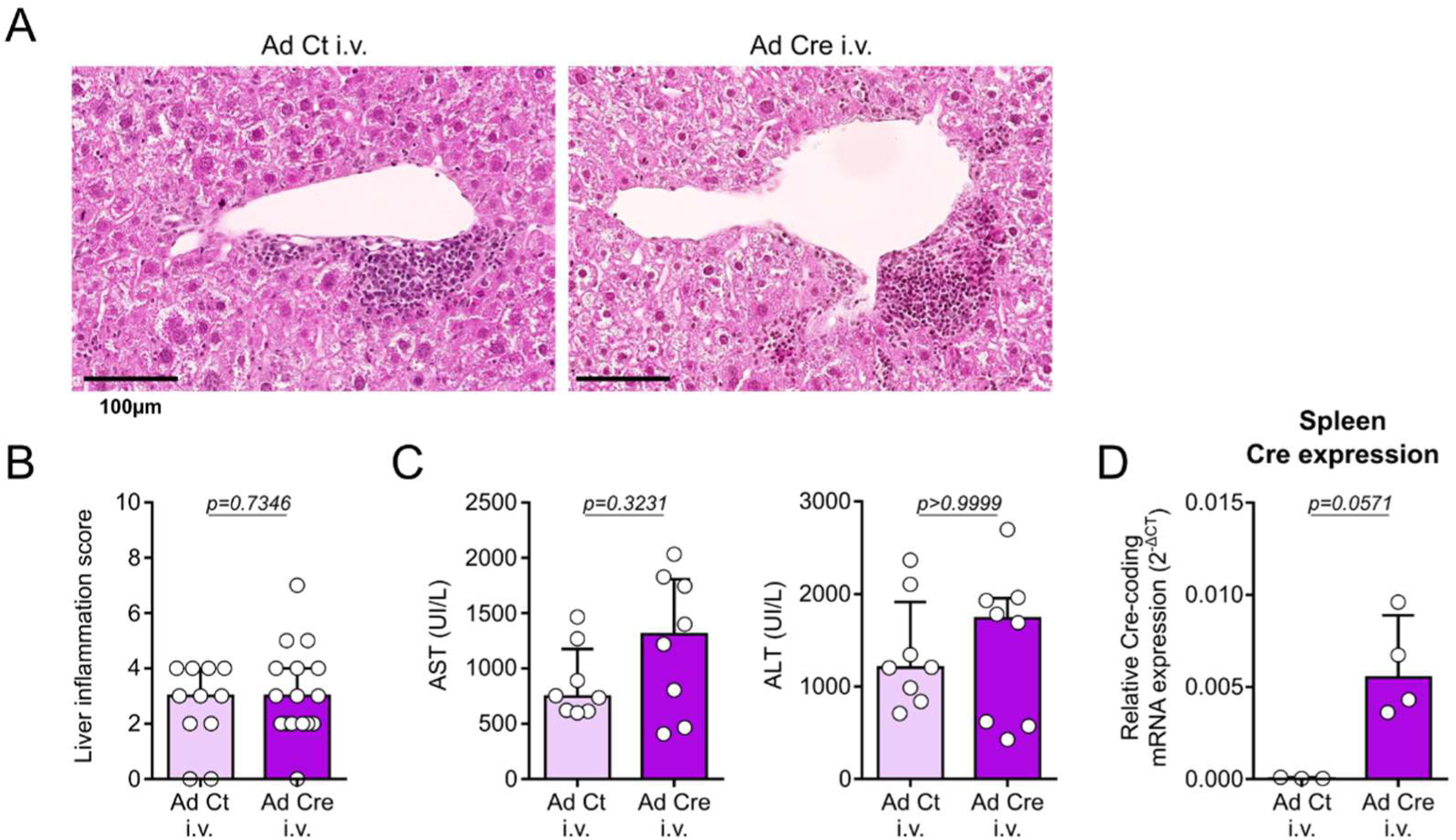
Intravenous injection of Ad Cre induces similar liver inflammation as Ad Ct i.v. and Cre expression in the spleen of mice. **(A)** Representative pictures of paraffin-embedded liver sections stained with HPS coloration from indicated conditions. **(B)** Histological liver inflammation scoring analysis of liver tissue sections from Ad Ct i.v. (n=11) and Ad Cre i.v. (n=16) mice. **(C)** Transaminase (AST/ALT) levels in serum of mice. **(D)** Relative Cre mRNA expression in the spleen of Ad Ct i.v. and Ad Cre i.v. mice. ACTB was used as loading control. Data are represented as median ± interquartile range in graphs B, C and D. Two-sided Mann-Whitney test was used for B, C and D. p-values are indicated.

**Supplementary Figure 2.**
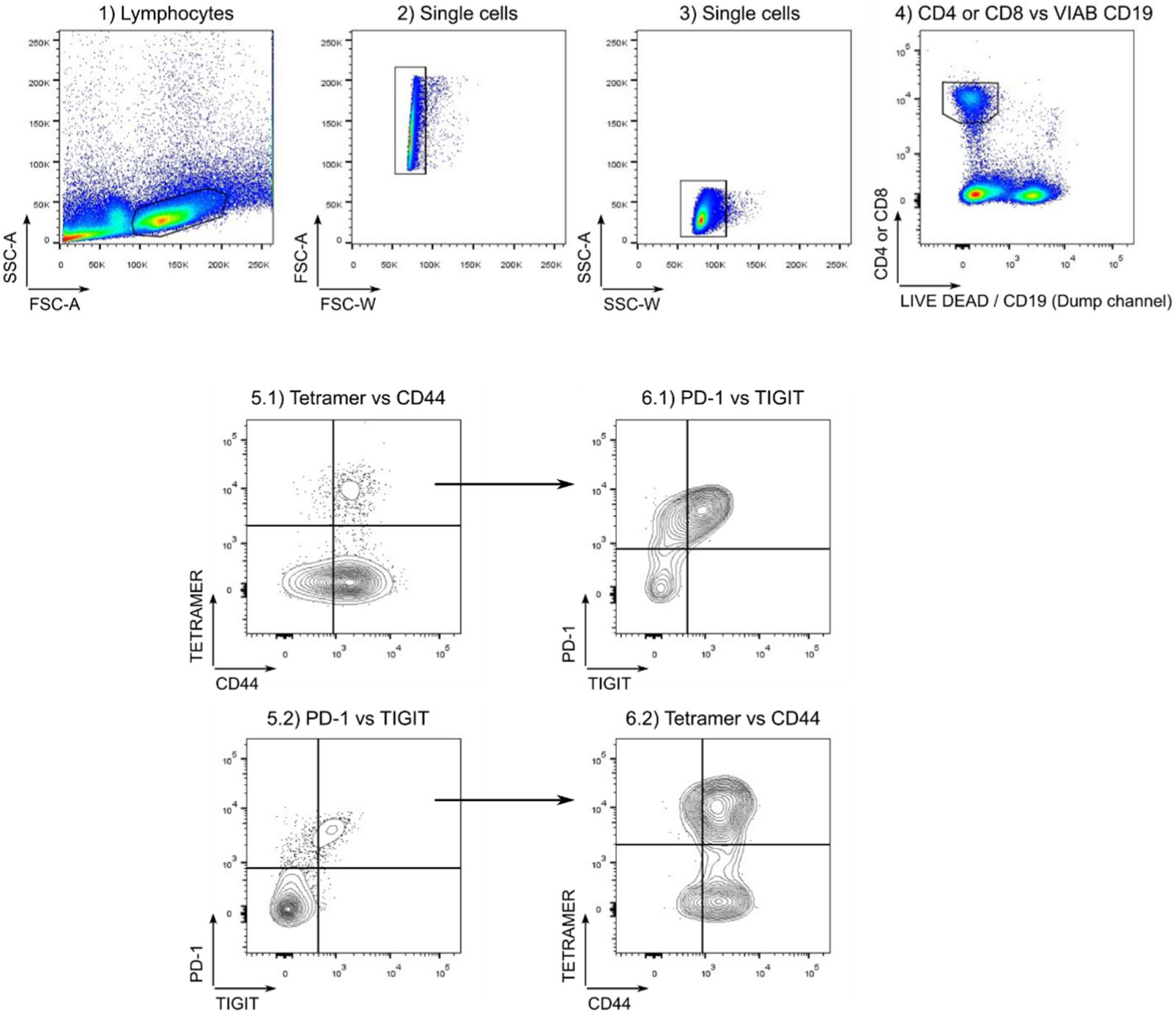
Conventional flow cytometry gating strategy for the analysis of HA-specific CD4 and CD8 T cells. HA-specific CD4 and CD8 T cells were defined as LIVE DEAD Aqua^-^ CD19^-^ CD4^+^/CD8^+^ CD44^high/+^ Tetramer^+^ cells. Data were acquired on BDFACSCanto.

**Supplementary Figure 3.**
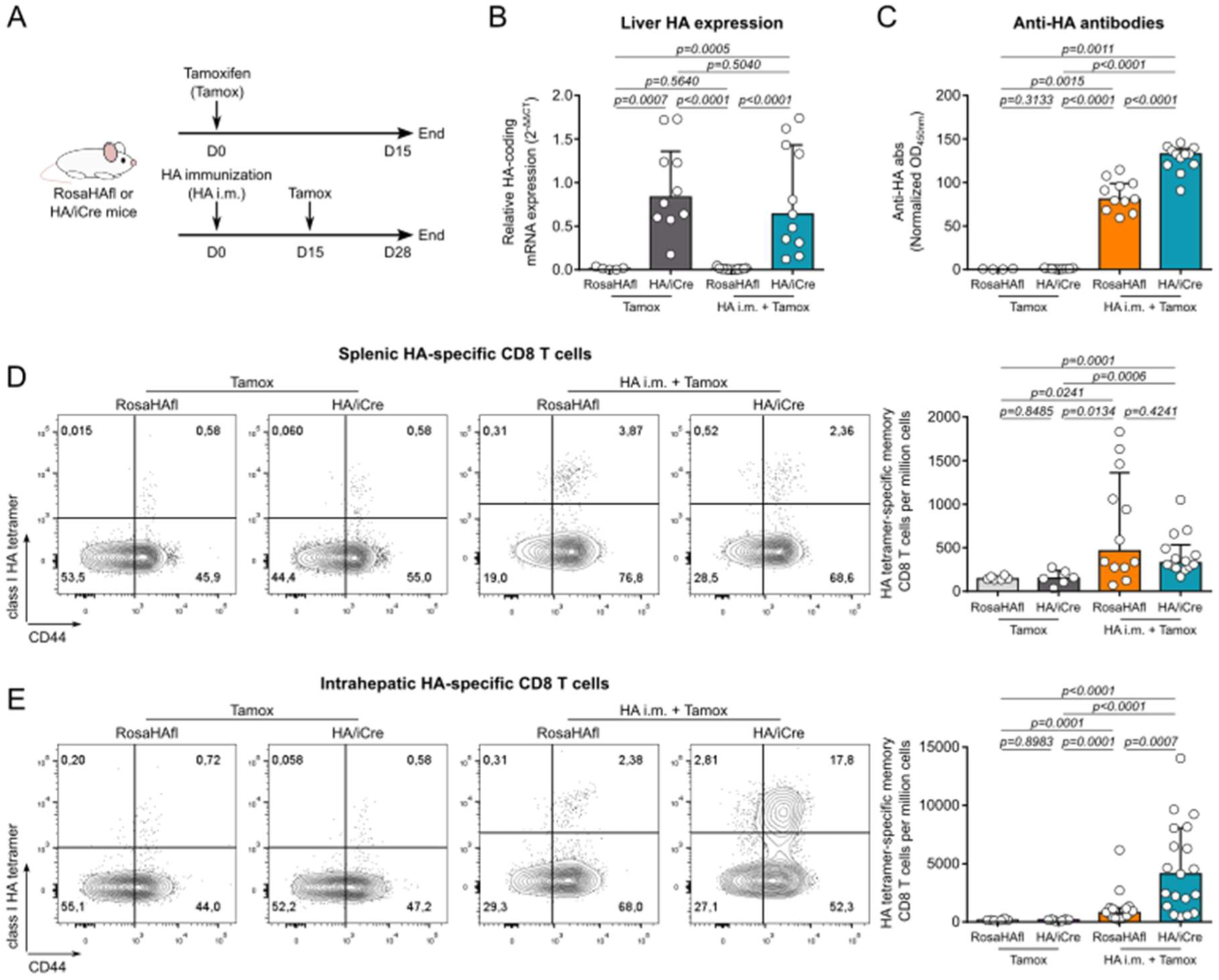
Tamoxifen treatment is not associated with a HA-specific CD8 T cell response in the spleen or in the liver. **(A)** Experimental design for the tamoxifen treatment in naive or HA immunized mice. RosaHAfl or HA/iCre mice received either tamoxifen treatment alone (Tamox, RosaHAfl, n=6; HA/iCre, n=10) or intramuscular HA immunization followed two weeks later by tamoxifen treatment (HA i.m. + Tamox, RosaHAfl, n=16; HA/iCre, n=19). Mice were euthanized two weeks after the last treatment. **(B)** Relative HA mRNA expression in the liver of mice. ACTB was used as loading control. **(C)** Normalized anti-HA antibody rate in serum of mice. (D-E) Analysis of HA- specific CD8 T cells in the spleen **(D)** and in the liver **(E)** of RosaHAfl and HA/iCre mice following Tamox or HA i.m. + Tamox protocols. Contour plot representation of HA tetramer staining and CD44 expression in CD8 T cells (left). Frequency of HA-specific memory (CD44^+^) CD8 T cells per million cells (right). Data are represented as median ± interquartile range in graphs B-E. Two-sided Mann- Whitney test was used for B-E. p-values are indicated.

**Supplementary Figure 4.**
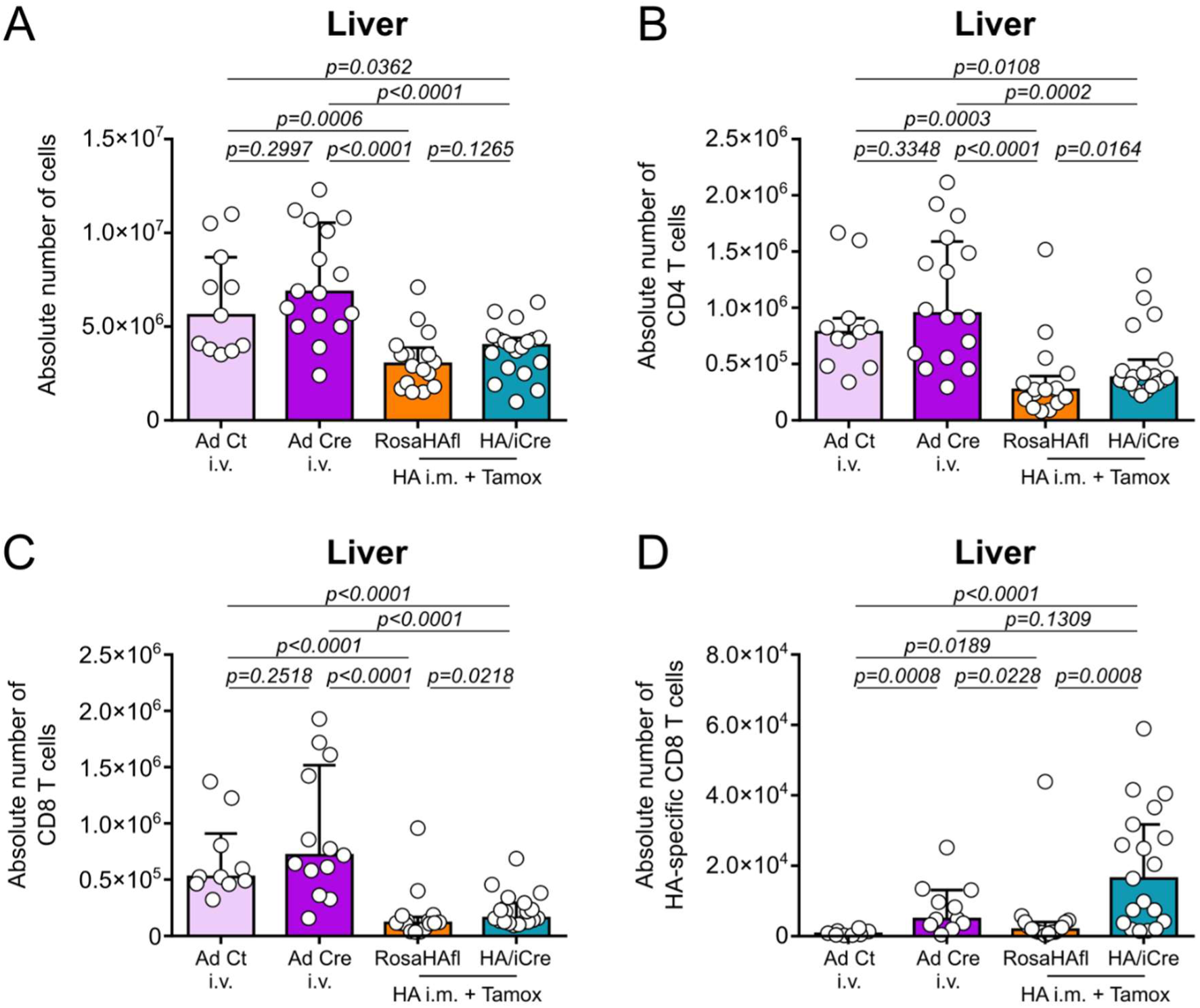
Intravenous injection of adenoviral vector induces high infiltration of T cells in the liver. **(A)**, CD4 T cells **(B)**, CD8 T cells **(C)** and HA-specific CD8 T cells **(D)** extracted from the liver of Ad Ct i.v. (n=11), Ad Cre i.v. (n=16), and RosaHAfl (n=16) or HA/iCre (n=19) mice that received HA i.m. + Tamox protocol. Data are represented as median ± interquartile range in graphs **A-D**. Two-sided Mann-Whitney test was used for **A-D**. p-values are indicated.

**Supplementary Figure 5.**
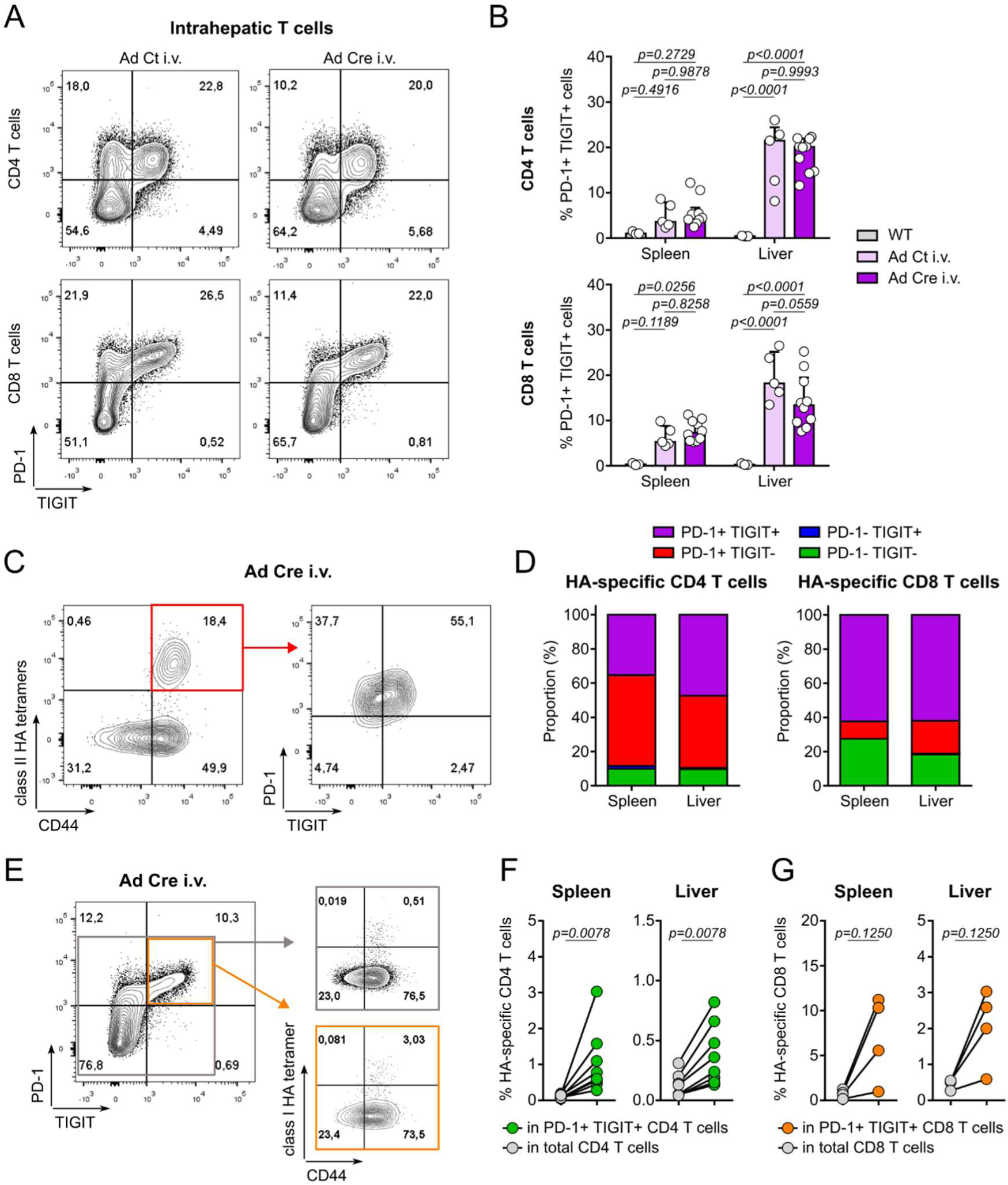
T cell reactivity towards adenoviral antigens is associated with PD-1 and TIGIT co-expression by T cells in the tissue. **(A)** Contour plot representation of PD-1 and TIGIT co-expression by CD4 T cells (top) and CD8 T cells (bottom) from the liver of Ad Ct i.v. (left) and Ad Cre i.v. (right) mice. **(B)** Analysis of PD-1/TIGIT co-expression by CD4 T cells (top) and CD8 T cells (bottom) from the spleen and the liver of wild-type Balb/c mice (n=3), Ad Ct i.v. (n=5) and Ad Cre i.v. (n=10) mice. **(C)** Contour plot representation of PD-1/TIGIT expression among HA-specific CD4 T cells (red). **(D)** Proportion of PD-1/TIGIT expressing subsets among HA-specific CD4 (left) and CD8 (right) T cells from the spleen and the liver of Ad Cre i.v. mice (n=8). **(E)** Contour plot representation of HA-specific CD8 T cell frequency among total CD8 T cells (grey) and PD-1^+^ TIGIT^+^ CD8 T cells (orange). **(F)** Analysis of HA-specific CD4 T cell frequency among total CD4 T cells and PD-1^+^ TIGIT^+^ CD4 T cells in the spleen and the liver. **(G)** Analysis of HA-specific CD8 T cell frequency among total CD8 T cells and PD-1^+^ TIGIT^+^ CD8 T cells in the spleen and the liver. Data are represented as mean in graph **D** and median ± interquartile range in graph **B.** Sidak’s multiple comparisons test was used for B. Wilcoxon matched-pairs signed rank test was used for **F** and **G**. p-values are indicated.

**Supplementary Figure 6.**
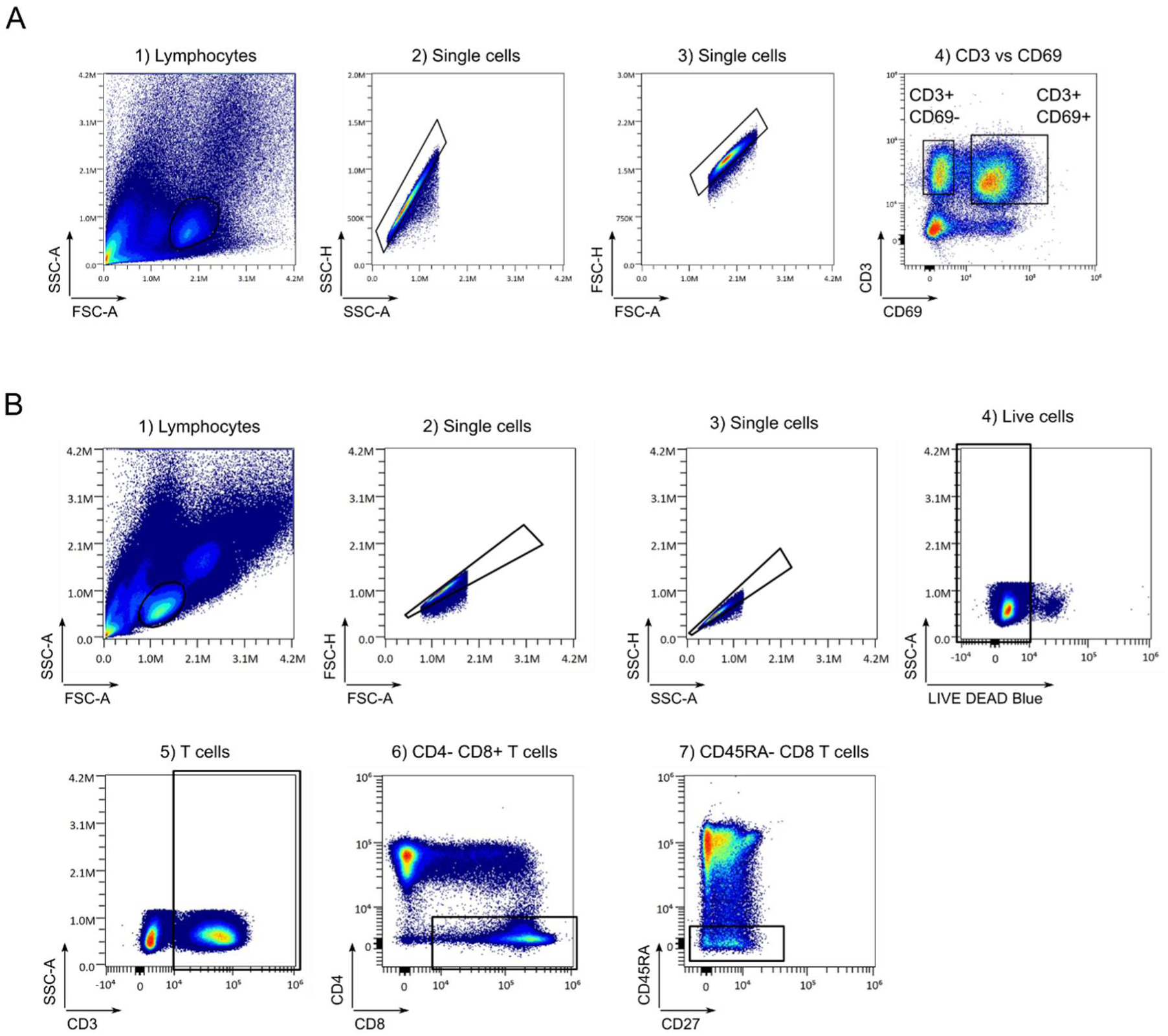
Spectral flow cytometry gating strategy for the analysis of intrahepatic T cells and blood memory CD8 T cells from AIH and NASH patients. **(A)** Pseudocolor dot plot representation of the gating strategy for CD69^-^ and CD69^+^ CD3^+^ cells on intrahepatic cells from AIH patient 10-004. **(B)** Pseudocolor dot plot representation of the gating strategy for CD45RA^-^ memory CD8 T cells from the blood of AIH and NASH patients. Data were acquired on Cytek Aurora flow cytometer.

**Supplementary Figure 7.**
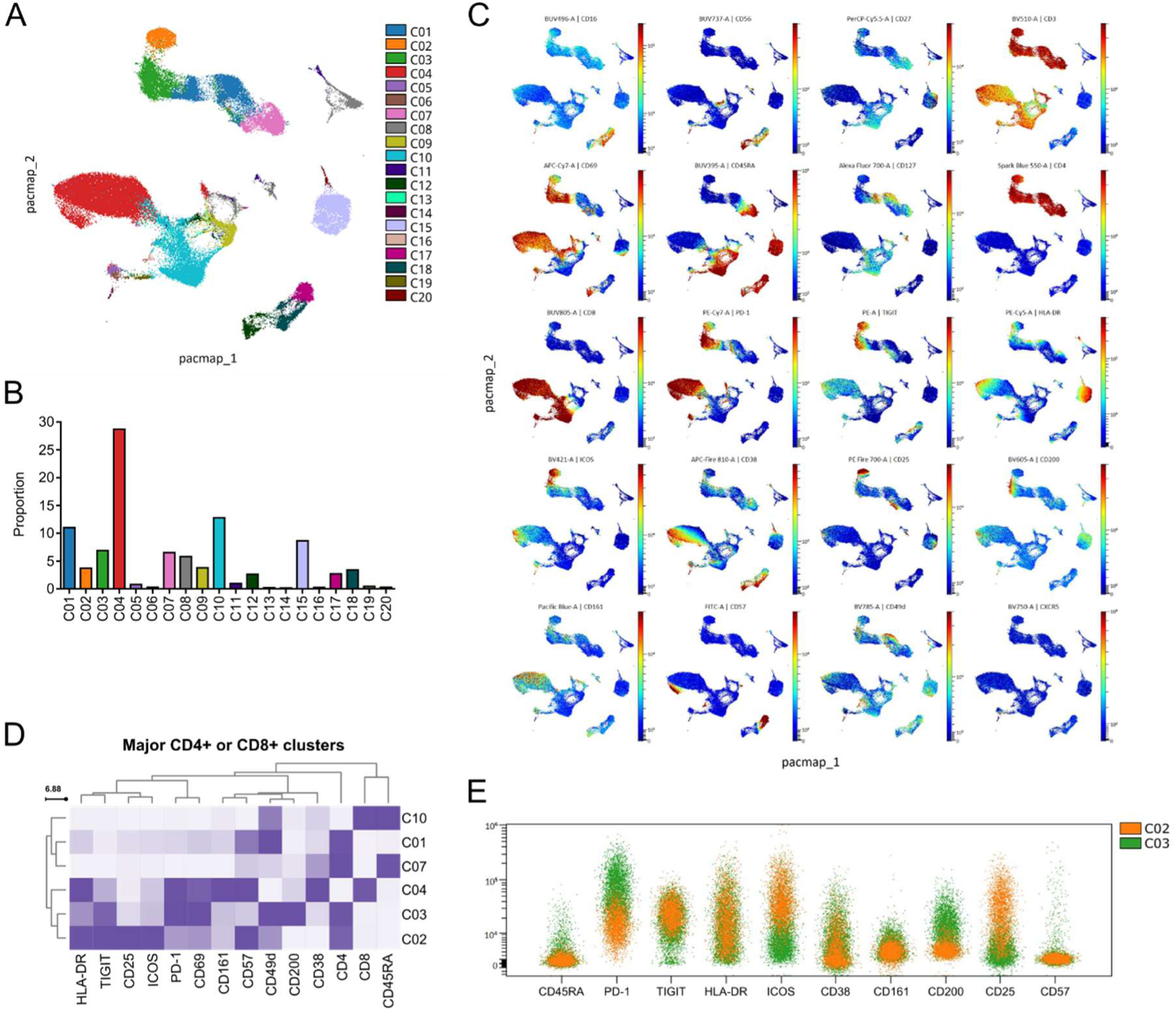
Unsupervised clustering analysis of intrahepatic T cells from AIH patient 10-004. **(A)** PaCMAP representation of intrahepatic cells from AIH patient 10-004, colored by unsupervised FlowSOM clustering. **(B)** Proportion of clusters defined in **A** among total intrahepatic cells. **(C)** Surface markers expression on PaCMAP representation defined in **A**. **(D)** Surface markers heatmap of major CD4+ (C02, C03, C07, C01) and CD8+ (C04, C10) clusters. **(E)** Overlaid spectrum plot of C02 and C03.

**Supplementary Figure 8.**
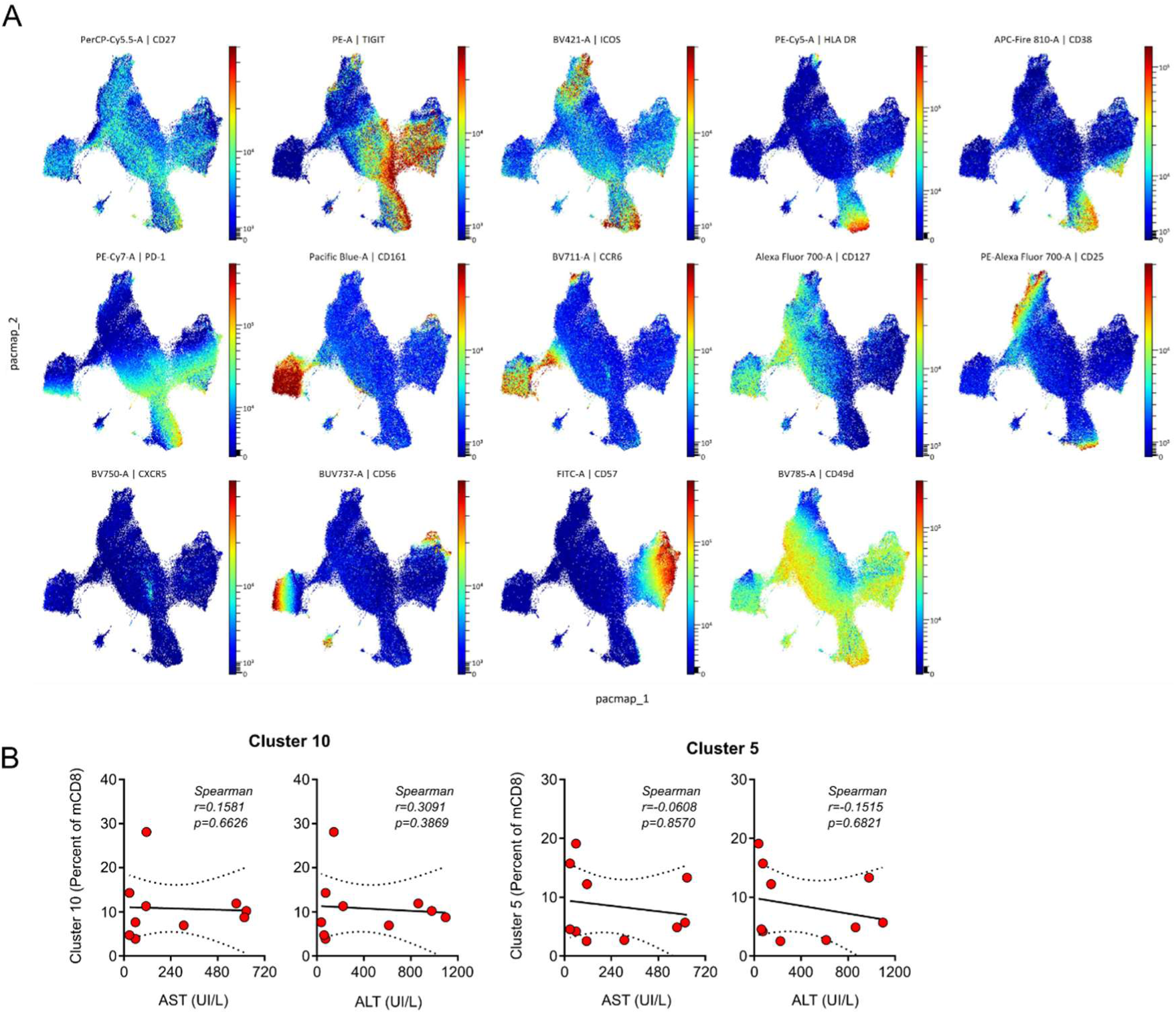
Frequency of PD-1^+^ TIGIT^+^ HLA-DR^-^ CD38^-^ ICOS^-^ memory CD8 T cells is not correlated with liver injury in active AIH patients. **(A)** Surface marker expression on PaCMAP representation of blood mCD8 from AIH and NASH patients defined in **Figure 7B**. **(B)** Correlation analysis of PD-1^+^ TIGIT^+^ HLA-DR^-^ CD38^-^ ICOS^-^ mCD8 C10 and C5 cluster frequencies among blood mCD8 with AST and ALT rate in AIHa patients. Spearman correlation test was used for **B**. p-values are indicated.

**Supplementary Figure 9.**
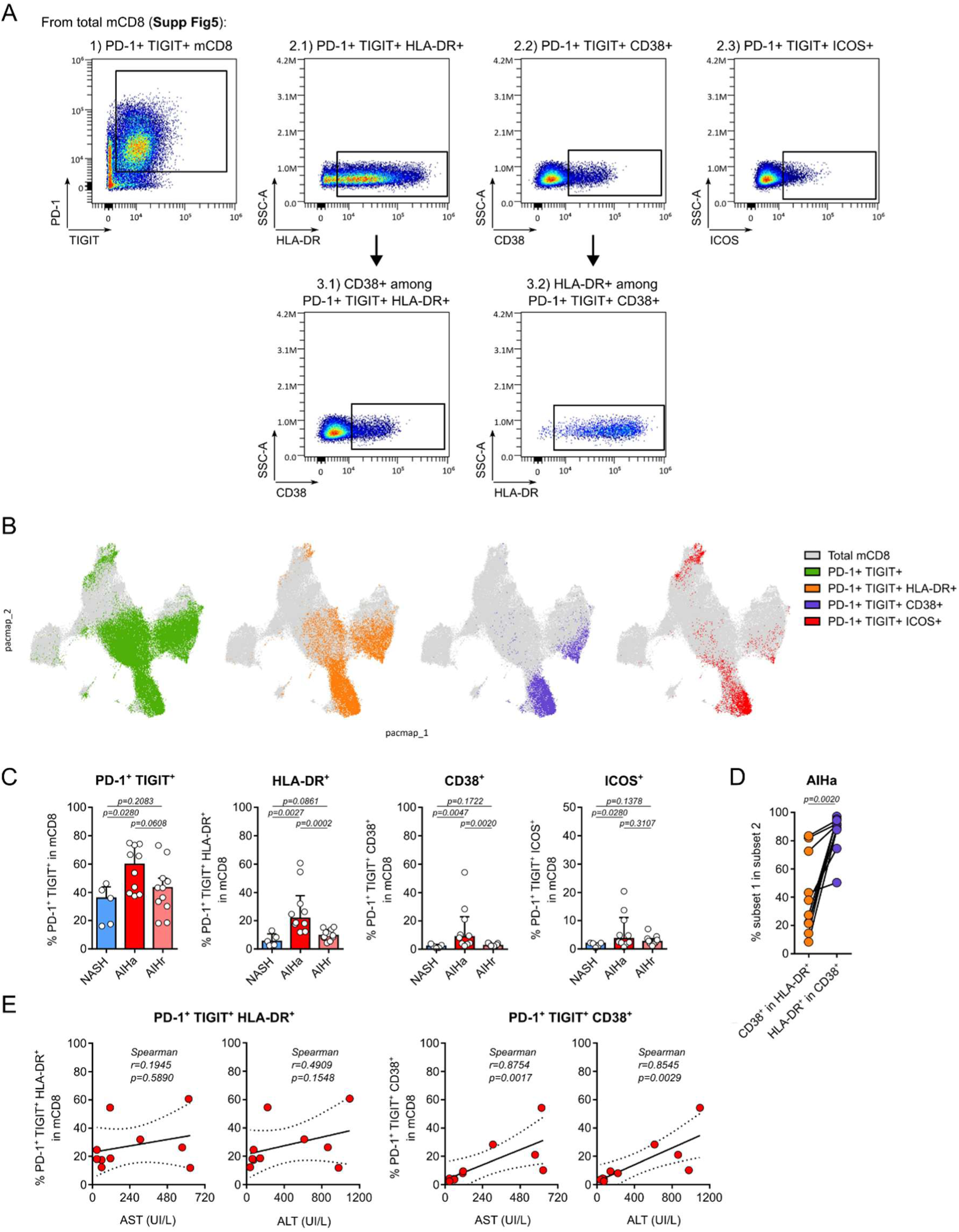
Frequency of circulating PD-1^+^ TIGIT^+^ CD38^+^ memory CD8 T cells correlates with liver injury in active AIH patients. **(A)** Pseudocolor dot plot representation of the supervised gating strategy for HLA-DR, CD38 and ICOS analysis in PD-1^+^ TIGIT^+^ memory CD8 T cells. **(B)** PaCMAP representation shown in **Figure 6B** colored by PD-1^+^ TIGIT^+^ mCD8 (green), PD-1^+^ TIGIT^+^ HLA-DR^+^ mCD8 (orange), PD-1^+^ TIGIT^+^ CD38^+^ mCD8 (purple) and PD-1^+^ TIGIT^+^ ICOS^+^ mCD8 (red). **(C)** Analysis of the frequency of subsets shown in B among mCD8 between NASH, AIHa and AIHr patients. **(D)** Analysis of CD38 and HLA-DR expression in PD-1^+^ TIGIT^+^ HLA-DR^+^ and CD38^+^ subsets, respectively, in AIHa patients. **(E)** Correlation analysis of the frequency of PD-1^+^ TIGIT^+^ HLA-DR^+^ and CD38^+^ among mCD8 with AST and ALT rate of AIHa patients. Data are represented as median ± interquartile range in graph **C**. Two-sided Mann-Whitney test was used for **C**. Wilcoxon matched-pairs signed rank test was used for **D**. Spearman correlation test was used for **E**. p-values are indicated.

**Supplementary Table 1.** Clinical and biological characteristics of patients with AIH and NASH expressed as mean [95% confidence interval].

| Group | Active AIH (AIHa) | Remission AIH (AIHr) | NASH |
| --- | --- | --- | --- |
|  | n=10 | n=11 | n=5 |
| Age (year) | 49,6 [37,7-61,5] | 49,5 [34,2-64,7] | 59,2 [33,3-85,1] |
| Female, n (%) | 6 (60%) | 9 (82%) | 3 (60%) |
| IgG (g/L) | 18,0 [13,5-22,5] | 11,1 [9,4-12,7] | 11,9 [6,8-17,1] |
| AST (IU/L) | 252,9 [69,2-436,5] | 21,2 [17,1-25,3] | 45,9 [23,4-68,4] |
| ALT (IU/L) | 416,6 [111,8-721,4] | 19,8 [12,7-26,8] | 73,5 [6,2-140,9] |
| Immunosuppressive agents n (%) |  |  |  |
| No, n (%) | 7 (70%) | 0 | 5 (100%) |
| Naïve | 7 | 0 | NA |
| Yes, n (%) | 3 (30%) | 11 (100%) | 0 |
| Azathioprin | 2 | 10 | NA |
| MMF | 1 | 0 | NA |
| Steroids only | 0 | 1 | NA |
| Steroids associated to other therapy | 2 | 2 | NA |

*AIH: Autoimmune Hepatitis, NASH: Non-Alcoholic Steatohepatitis, AST: Aspartate Aminotransferase,*
*ALT: Alanine Aminotransferase, NA: non-applicable*

**Supplementary Table 2.**
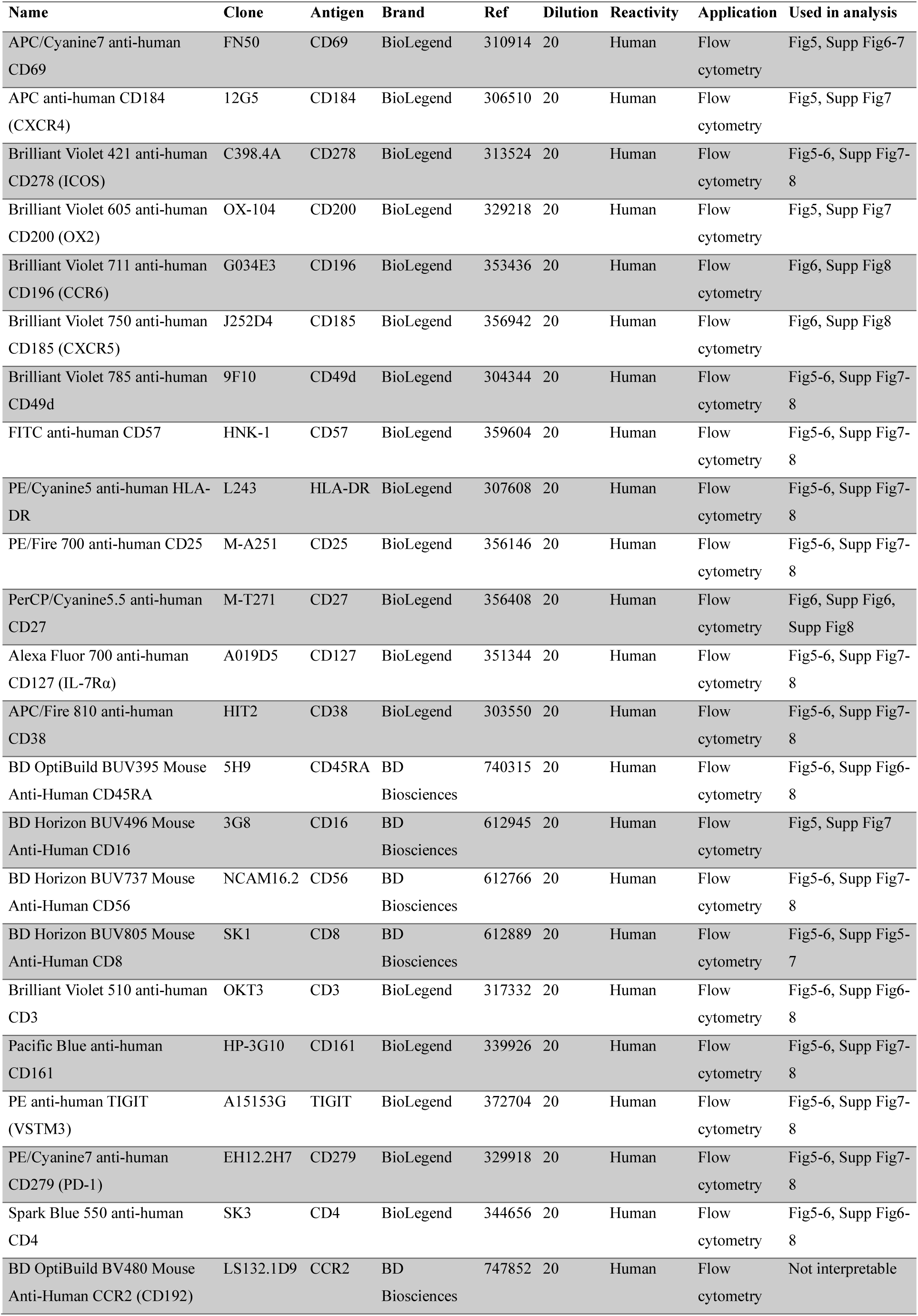
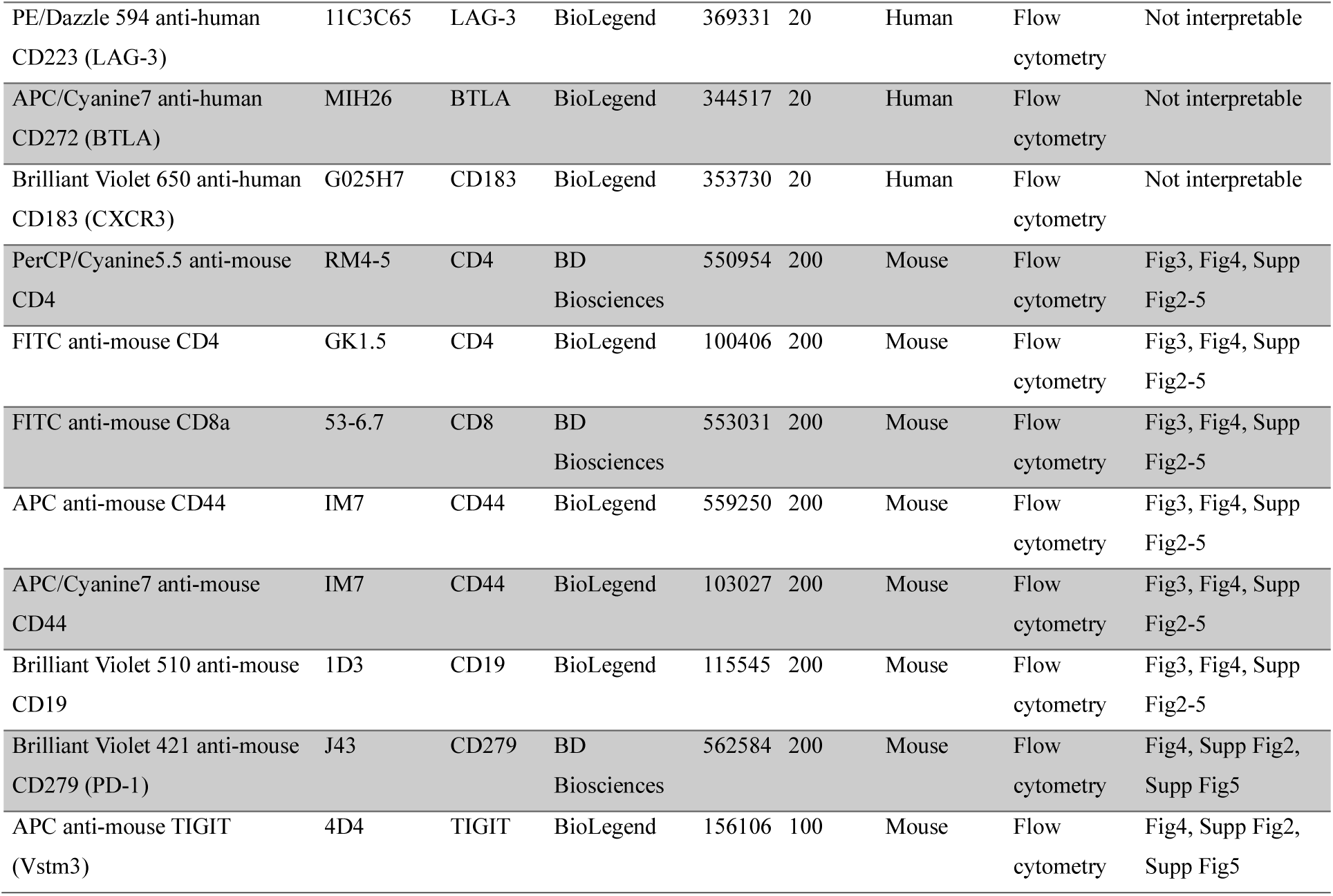
List of labeled antibodies.

| Name | Clone | Antigen | Brand | Ref | Dilution | Reactivity | Application | Used in analysis |
| --- | --- | --- | --- | --- | --- | --- | --- | --- |
| APC/Cyanine7 anti-human CD69 | FN50 | CD69 | BioLegend | 310914 | 20 | Human | Flow cytometry | Fig5, Supp Fig6-7 |
| APC anti-human CD184 (CXCR4) | 12G5 | CD184 | BioLegend | 306510 | 20 | Human | Flow cytometry | Fig5, Supp Fig7 |
| Brilliant Violet 421 anti-human CD278 (ICOS) | C398.4A | CD278 | BioLegend | 313524 | 20 | Human | Flow cytometry | Fig5-6, Supp Fig7-8 |
| Brilliant Violet 605 anti-human CD200 (OX2) | OX-104 | CD200 | BioLegend | 329218 | 20 | Human | Flow cytometry | Fig5, Supp Fig7 |
| Brilliant Violet 711 anti-human CD196 (CCR6) | G034E3 | CD196 | BioLegend | 353436 | 20 | Human | Flow cytometry | Fig6, Supp Fig8 |
| Brilliant Violet 750 anti-human CD185 (CXCR5) | J252D4 | CD185 | BioLegend | 356942 | 20 | Human | Flow cytometry | Fig6, Supp Fig8 |
| Brilliant Violet 785 anti-human CD49d | 9F10 | CD49d | BioLegend | 304344 | 20 | Human | Flow cytometry | Fig5-6, Supp Fig7-8 |
| FITC anti-human CD57 | HNK-1 | CD57 | BioLegend | 359604 | 20 | Human | Flow cytometry | Fig5-6, Supp Fig7-8 |
| PE/Cyanine5 anti-human HLA-DR | L243 | HLA-DR | BioLegend | 307608 | 20 | Human | Flow cytometry | Fig5-6, Supp Fig7-8 |
| PE/Fire 700 anti-human CD25 | M-A251 | CD25 | BioLegend | 356146 | 20 | Human | Flow cytometry | Fig5-6, Supp Fig7-8 |
| PerCP/Cyanine5.5 anti-human CD27 | M-T271 | CD27 | BioLegend | 356408 | 20 | Human | Flow cytometry | Fig6, Supp Fig6, Supp Fig8 |
| Alexa Fluor 700 anti-human CD127 (IL-7Rα) | A019D5 | CD127 | BioLegend | 351344 | 20 | Human | Flow cytometry | Fig5-6, Supp Fig7-8 |
| APC/Fire 810 anti-human CD38 | HIT2 | CD38 | BioLegend | 303550 | 20 | Human | Flow cytometry | Fig5-6, Supp Fig7-8 |
| BD OptiBuild BUV395 Mouse Anti-Human CD45RA | 5H9 | CD45RA | BD Biosciences | 740315 | 20 | Human | Flow cytometry | Fig5-6, Supp Fig6-8 |
| BD Horizon BUV496 Mouse Anti-Human CD16 | 3G8 | CD16 | BD Biosciences | 612945 | 20 | Human | Flow cytometry | Fig5, Supp Fig7 |
| BD Horizon BUV737 Mouse Anti-Human CD56 | NCAM16.2 | CD56 | BD Biosciences | 612766 | 20 | Human | Flow cytometry | Fig5-6, Supp Fig7-8 |
| BD Horizon BUV805 Mouse Anti-Human CD8 | SK1 | CD8 | BD Biosciences | 612889 | 20 | Human | Flow cytometry | Fig5-6, Supp Fig5-7 |
| Brilliant Violet 510 anti-human CD3 | OKT3 | CD3 | BioLegend | 317332 | 20 | Human | Flow cytometry | Fig5-6, Supp Fig6-8 |
| Pacific Blue anti-human CD161 | HP-3G10 | CD161 | BioLegend | 339926 | 20 | Human | Flow cytometry | Fig5-6, Supp Fig7-8 |
| PE anti-human TIGIT (VSTM3) | A15153G | TIGIT | BioLegend | 372704 | 20 | Human | Flow cytometry | Fig5-6, Supp Fig7-8 |
| PE/Cyanine7 anti-human CD279 (PD-1) | EH12.2H7 | CD279 | BioLegend | 329918 | 20 | Human | Flow cytometry | Fig5-6, Supp Fig7-8 |
| Spark Blue 550 anti-human CD4 | SK3 | CD4 | BioLegend | 344656 | 20 | Human | Flow cytometry | Fig5-6, Supp Fig6-8 |
| BD OptiBuild BV480 Mouse Anti-Human CCR2 (CD192) | LS132.1D9 | CCR2 | BD Biosciences | 747852 | 20 | Human | Flow cytometry | Not interpretable |
| PE/Dazzle 594 anti-human CD223 (LAG-3) | 11C3C65 | LAG-3 | BioLegend | 369331 | 20 | Human | Flow cytometry | Not interpretable |
| APC/Cyanine7 anti-human CD272 (BTLA) | MIH26 | BTLA | BioLegend | 344517 | 20 | Human | Flow cytometry | Not interpretable |
| Brilliant Violet 650 anti-human CD183 (CXCR3) | G025H7 | CD183 | BioLegend | 353730 | 20 | Human | Flow cytometry | Not interpretable |
| PerCP/Cyanine5.5 anti-mouse CD4 | RM4-5 | CD4 | BD Biosciences | 550954 | 200 | Mouse | Flow cytometry | Fig3, Fig4, Supp Fig2-5 |
| FITC anti-mouse CD4 | GK1.5 | CD4 | BioLegend | 100406 | 200 | Mouse | Flow cytometry | Fig3, Fig4, Supp Fig2-5 |
| FITC anti-mouse CD8a | 53-6.7 | CD8 | BD Biosciences | 553031 | 200 | Mouse | Flow cytometry | Fig3, Fig4, Supp Fig2-5 |
| APC anti-mouse CD44 | IM7 | CD44 | BioLegend | 559250 | 200 | Mouse | Flow cytometry | Fig3, Fig4, Supp Fig2-5 |
| APC/Cyanine7 anti-mouse CD44 | IM7 | CD44 | BioLegend | 103027 | 200 | Mouse | Flow cytometry | Fig3, Fig4, Supp Fig2-5 |
| Brilliant Violet 510 anti-mouse CD19 | 1D3 | CD19 | BioLegend | 115545 | 200 | Mouse | Flow cytometry | Fig3, Fig4, Supp Fig2-5 |
| Brilliant Violet 421 anti-mouse CD279 (PD-1) | J43 | CD279 | BD Biosciences | 562584 | 200 | Mouse | Flow cytometry | Fig4, Supp Fig2, Supp Fig5 |
| APC anti-mouse TIGIT (Vstm3) | 4D4 | TIGIT | BioLegend | 156106 | 100 | Mouse | Flow cytometry | Fig4, Supp Fig2, Supp Fig5 |

**Supplementary Table 3.**
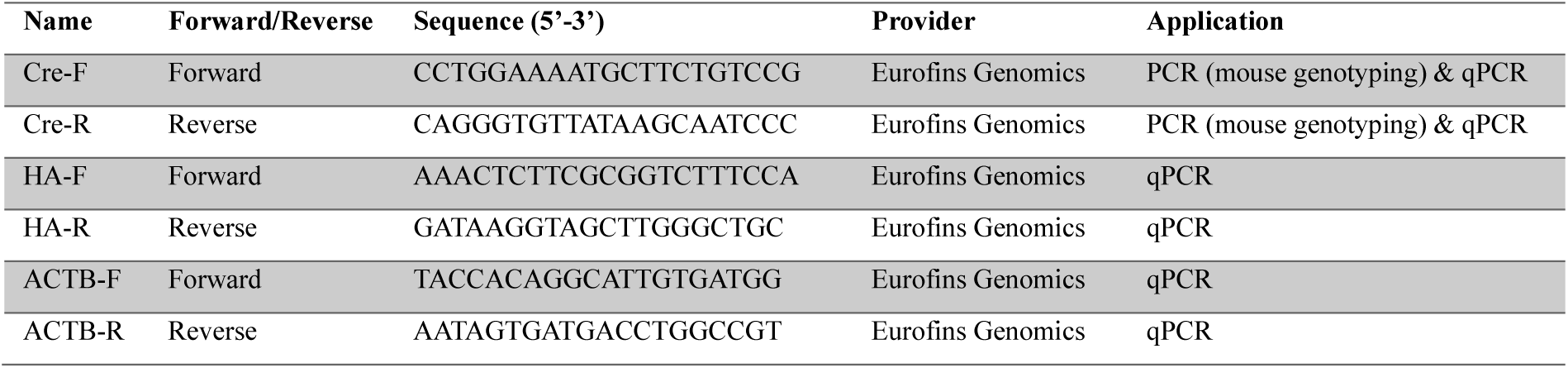
PCR and qPCR primers.

